# On the cusp of adaptive change: the hierarchical radiation of phyllostomid bats

**DOI:** 10.1101/2023.05.23.541856

**Authors:** David M. Grossnickle, Alexa Sadier, Edward Patterson, Nashaly N. Cortés-Viruet, Stephanie Jimenez Rivera, Karen E. Sears, Sharlene E. Santana

## Abstract

Adaptive radiations are bursts in biodiversity that lead to the origin of new evolutionary lineages and phenotypes. However, adaptive radiations typically occur over millions of years and it is unclear how the macroevolutionary dynamics that underpin them vary through time and among groups of organisms. Phyllostomid bats radiated extensively for diverse diets –from insects to vertebrates, fruit, nectar, and blood– and we use their molars as a model system to examine the dynamics of adaptive radiations. Three-dimensional shape analyses of lower molars of Noctilionoidea (Phyllostomidae and close relatives) indicate that different diet groups exhibit distinct morphotypes. Comparative analyses further reveal that phyllostomids are a striking example of a hierarchical radiation; their initial, higher-level diversification involved an ‘early burst’ in molar morphological disparity as lineages invaded new diet-affiliated adaptive zones, followed by subsequent lower-level diversifications within adaptive zones involving less dramatic morphological changes. We posit that strong selective pressures related to initial shifts to derived diets may have freed molars from morpho-functional constraints associated with the ancestral molar morphotype. Then, lineages with derived diets (frugivores and nectarivores) diversified considerably within broad adaptive zones, likely reflecting finer-scale niche partitioning. The observed early burst pattern is only evident when examining molar traits that are strongly linked to diet, highlighting the importance of ecomorphological traits in comparative studies. Our results support the hypothesis that adaptive radiations are commonly hierarchical and involve different tempos and modes at different phylogenetic scales, with early bursts being more common at broader scales.

**SIGNIFICANCE STATEMENT:** Many groups of organisms are exceptionally diverse in their ecology, morphology, and number of species. But there is debate as to whether these groups commonly achieved this diversity through ‘bursts’ in diversification early in their history. Phyllostomid bats are one of the most ecologically diverse mammalian families and a classic example of an adaptive radiation. We use their molar shapes, which correlate with diet, as a model for examining macroevolutionary patterns during diversifications. We find that phyllostomids experienced a two-step process of diversification; the first step involved a rapid burst, whereas the second involved finer-scale changes as lineages filled ecological niches. We posit that this is a common, yet underappreciated, pattern during the early histories of many diverse clades.

## INTRODUCTION

Novel biodiversity is often the product of adaptive radiations, which occur in response to new ecological opportunities and are marked by rapid increases in ecological, morphological, and taxonomic diversity (e.g., Osborn 1902, Simpson 1944, Erwin 1992, Schluter 2000, Givnish 2015, Stroud and Losos 2016).

Nonetheless, there remains considerable uncertainty about the macroevolutionary dynamics that characterize adaptive radiations. An oft-hypothesized pattern of adaptive radiations is an ‘early burst’ in ecomorphological change as lineages rapidly adapt to new niches, followed by a slowing of evolutionary rates as ecological opportunity diminishes (Simpson 1944, Simpson 1953, Harmon et al. 2003, Gavrilets and Losos 2009, Harmon et al. 2010). Simpson’s (1944) seminal conceptualization of this pattern involved a rapid invasion of new adaptive zones by lineages (i.e., an ‘explosive phase’ involving ‘quantum evolution’), followed by finer-scale niche partitioning within adaptive zones. But despite the appeal and popularity of this hypothesis, the early burst pattern is rarely observed in phylogenetic comparative analyses, especially those examining extant-only samples (Harmon et al. 2010, Slater et al. 2012, Mitchell 2015, Pincheira-Donoso et al. 2015, Arbour et al. 2019, Grossnickle 2020, Weaver and Grossnickle 2020; but see counterexamples such as Arbour and López-Fernández 2013, Price et al. 2016, Stanchak et al. 2019). The fossil record provides some support for the early burst hypothesis, with clades often experiencing their greatest morphological diversity early in their history (Foote 1994, Ruta et al. 2006, Erwin 2007, Hughes et al. 2013, Grossnickle et al. 2019, Zhang and Shu 2021) and/or experiencing diversifications posited to be linked to new ecological opportunities (e.g., Simpson 1944, Wilson et al. 2012, Grossnickle and Newham 2016, Halliday and Goswami 2016). However, it is challenging to verify if these observed fossil diversifications represent adaptive radiations because the available evidence is fragmentary and ecological breadth is often difficult to infer in extinct taxa; in many cases, they could instead reflect ‘non-adaptive radiations’ (e.g., Erwin 2015, Givnish 2015). The dearth of quantifiable early bursts has led some researchers to challenge the hypothesis that early bursts are a common signature of adaptive radiations (e.g., Harmon et al. 2010, Pincheira-Donoso et al. 2015).

The ability to detect early bursts, however, may be hindered by multiple factors that bias the results of evolutionary model-fitting analyses, an oft-used comparative tool for examining macroevolutionary patterns. For instance, the fit of an early burst (EB) model (Harmon et al. 2010) to morphological datasets may be weakened by a lack of fossil taxa (e.g., Slater et al. 2012, Mitchell 2015), small sample size, and convergent evolution among lineages of different adaptive zones (Slater and Pennell 2014). The choice of traits to examine also heavily influences model-fitting results. Because the early burst concept is associated with major ecological shifts, ecomorphological traits (e.g., tooth traits that reflect diet) are the most appropriate morphological data for testing early patterns in adaptive radiations (Schluter 2000, Colombo et al. 2015, Feilich and López-Fernández 2019, Slater and Friscia 2019, Grossnickle 2020, Slater 2022), but these are not always used or available. Thus, even in focal clades that experienced early bursts, researchers may fail to observe the pattern due to methodological choices.

The phylogenetic scale of the examined radiation is an additional factor that can influence the ability to detect early bursts. Based on Simpson’s (1944, 1953) model of adaptive radiation, an early burst pattern is *only* expected in higher-level radiations (‘general radiations’ of Osborn 1902) involving the appearance of new ecomorphotypes and invasion of new adaptive zones (Slater and Friscia 2019), such as taxa transitioning to derived diets or locomotor modes. In contrast to higher-level radiations, an early burst is expected to be less common in lower-level, within-adaptive-zone radiations (e.g., those at the subfamily or genus level; ‘local radiations’ of Osborn 1902), which could involve finer-scale resource and microhabitat partitioning (Simpson 1944). In support of these expectations, Slater and Friscia (2019) found that ecomorphological traits associated with diet in carnivorans demonstrate an early burst at the order level, but this pattern is less common at the family level. However, to the best of our knowledge, this pattern of hierarchical radiation has not been clearly demonstrated in additional taxa.

We use Neotropical leaf-nosed bats, family Phyllostomidae, and their close relatives (Noctilionoidea) as a model system for testing the hierarchical adaptive radiation hypothesis, which is characterized by an initial early-burst phase of evolution in ecologically-relevant traits. The early diversification of phyllostomids is often categorized as an adaptive radiation that is posited to have been driven by novel ecological opportunities coupled with strong selective pressures on craniodental features that enhanced feeding performance on novel diets (Dávalos 2007, Rojas et al. 2011, Dumont et al. 2012, Rossoni et al. 2017, Rossoni et al. 2019, Fleming et al. 2020). Starting approximately 30 million years ago, insectivorous or omnivorous phyllostomid lineages rapidly diversified to frugivory, sanguivory, nectarivory, and vertivory (Freeman 2000, Rojas et al. 2011, Baker et al. 2012, Gunnell et al. 2014, Hall et al. 2021). This dietary diversification is reflected in their extensive craniodental adaptations, such as those of the skull, jaw musculature, molars, and sensory systems (Freeman 2000, Monteiro and Nogueira 2010, Monteiro and Nogueira 2011, Santana et al. 2010, Santana et al. 2011, Santana et al. 2012, Dumont et al. 2014, Hedrick and Dumont 2018, Sadier et al. 2018, Fleming et al. 2020, Hall et al. 2021, Hedrick et al. 2021, Shi et al. 2021, López-Aguirre et al. 2022).

We apply comparative methods to a comprehensive three-dimensional molar shape dataset of noctilionoids. Molar shape and diet are strongly linked in phyllostomids and mammals in general, and thus provide us with ideal ecomorphological traits for examining adaptive radiations (Feilich and López-Fernández 2019, Slater and Friscia 2019, Grossnickle 2020, Slater 2022, López-Aguirre et al. 2022, Villalobos-Chaves and Santana 2022). We find strong support for the hierarchical adaptive radiation hypothesis (Osborn 1902, Simpson 1944, Simpson 1953, Slater and Friscia 2019); phyllostomids only show evidence of an early burst in molar morphology at higher phylogenetic scales that include lineages that evolved into new adaptive zones associated with derived diets. Further, we find no evidence of an early burst when using morphological traits that are not strongly linked to major diet categories. Thus, phyllostomids are a striking example of a hierarchical adaptive radiation, but this pattern is only apparent at broad phylogenetic scales and when using ecologically-relevant morphological data.

## RESULTS

### Molar correlates of diet

Noctilionoid molars occupy three distinct regions in PC1 and PC2 morphospace, corresponding closely to faunivory, frugivory, and nectarivory (Figs. 1C and S1). Sanguivores (vampire bats) would likely occupy a fourth region of morphospace, but they were excluded from our analyses because their first lower molar are too derived for collection of landmarks for geometric morphometrics (Fig. S2).

**Fig. 1.**
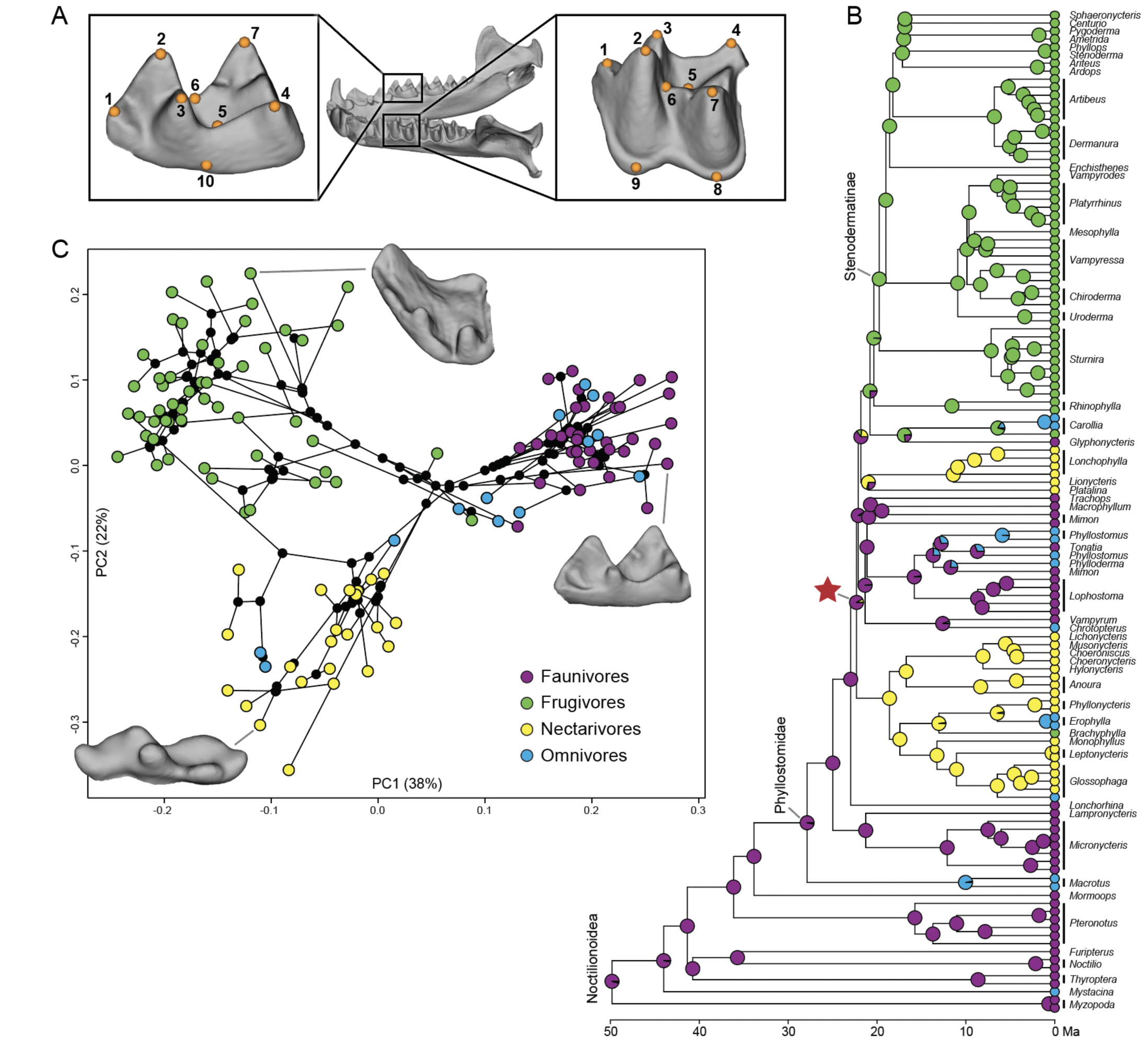
The molar landmarking scheme (*A*) and phylogeny (*B*) used in this study, with diets at ancestral nodes being summaries from 100 SIMMAP reconstructions. The 10 molar landmarks are described in the Supplementary Methods (SI Appendix). The PC1 and PC2 phylomorphospace plot (*C*) highlights the distinct ecomorphotypes associated with faunivory, frugivory, and nectarivory. See Figure S1 for a PCA plot with species labeled. The specimen images are of *Lophostoma silvicolum* (*A*; AMNH 64029), *Sphaeronycteris toxophyllum* (frugivore; AMNH 262637), *Lampronycteris brachyotis* (faunivore; AMNH 175639), and *Musonycteris harrisoni* (nectarivore; UMMZ 110524). The red star marks the node at which the first non-faunivore/omnivore lineages arise (“Phyllostomidae*”, see text), and this node is highlighted in subsequent figures.

We describe PC1–2 as ‘molar correlates of diet’ and use these PCs exclusively in some iterations of our analyses. Our decision to use these two PCs is based largely on results of pANOVAs and pMANOVAs, which demonstrate considerable molar shape differences among dietary groups in PC1-2 (*p* < 0.0001 for both diet schemes; Table S1). In contrast, all other PCs (PC3–30) and basicranium width (a proxy for body size) show no statistically significant differences among dietary groups in pANOVAs.

Functional considerations also support our decision to use PC1–2 as molar correlates of diet. PC1 (37% of variance) largely captures heights of molar cusps relative to overall tooth height, with faunivores (positive PC1 scores) having cusps that are relatively taller than those of frugivores and nectarivores (negative PC1 scores). Relative cusp heights are relevant functional specializations to dietary demands in bats, as demonstrated by the strong association between the molar relief index (the ratio of 3D surface area to 2D surface area) and diet (López-Aguirre et al. 2022, Villalobos-Chaves and Santana 2022), which is largely driven by differences in cusp heights among major diet groups (Boyer et al. 2008). PC2 (23% of variance) reflects relative differences in the lengths and widths of the molar, with frugivore molars (positive PC2 scores) being relatively wide and nectarivore molars (negative PC2 scores) being relatively long. Frugivores’ relatively wider molars may reflect a relatively greater occlusal area (López-Aguirre et al. 2022), which likely assists in crushing or grinding of seeds and fruit pulp. Finally, only PC1 and PC2 account for more variance than expected by chance via a ‘broken-stick’ model (Fig. S3; Jackson 1993), suggesting that these two PCs are capturing the most functionally relevant data.

### Morphological disparity patterns

The morphological-disparity-through-time patterns vary depending on the molar shape data examined (Fig. 2, Table 1). When using only molar correlates of diet (PC1–2), there is a distinct decrease in average subclade disparity at the node that we refer to as “Phyllostomidae*,” which is where Lonchorhinini + Micronycterini + Macrotinae splits from the rest of Phyllostomidae (red star in Figures 1B and 2A) and where most of Phyllostomidae’s dietary diversity begins to arise (Fig. 2A; see blue arrow). The morphological disparity index (MDI) results indicate that the average subclade disparity is lower than expected by chance for the major focal nodes in this study (Noctilionoidea, Phyllostomidae, and Phyllostomidae*; Table 1), which is consistent with an early burst pattern (Harmon et al. 2003, Slater et al. 2010). However, the distinct decrease in subclade disparity at the Phyllostomidae* node disappears when including molar shape data unrelated to diet (PC1–30; Fig. 2B), and subclades instead show elevated disparity until the Present (blue arrow in Figure 2B).

**Fig. 2.**
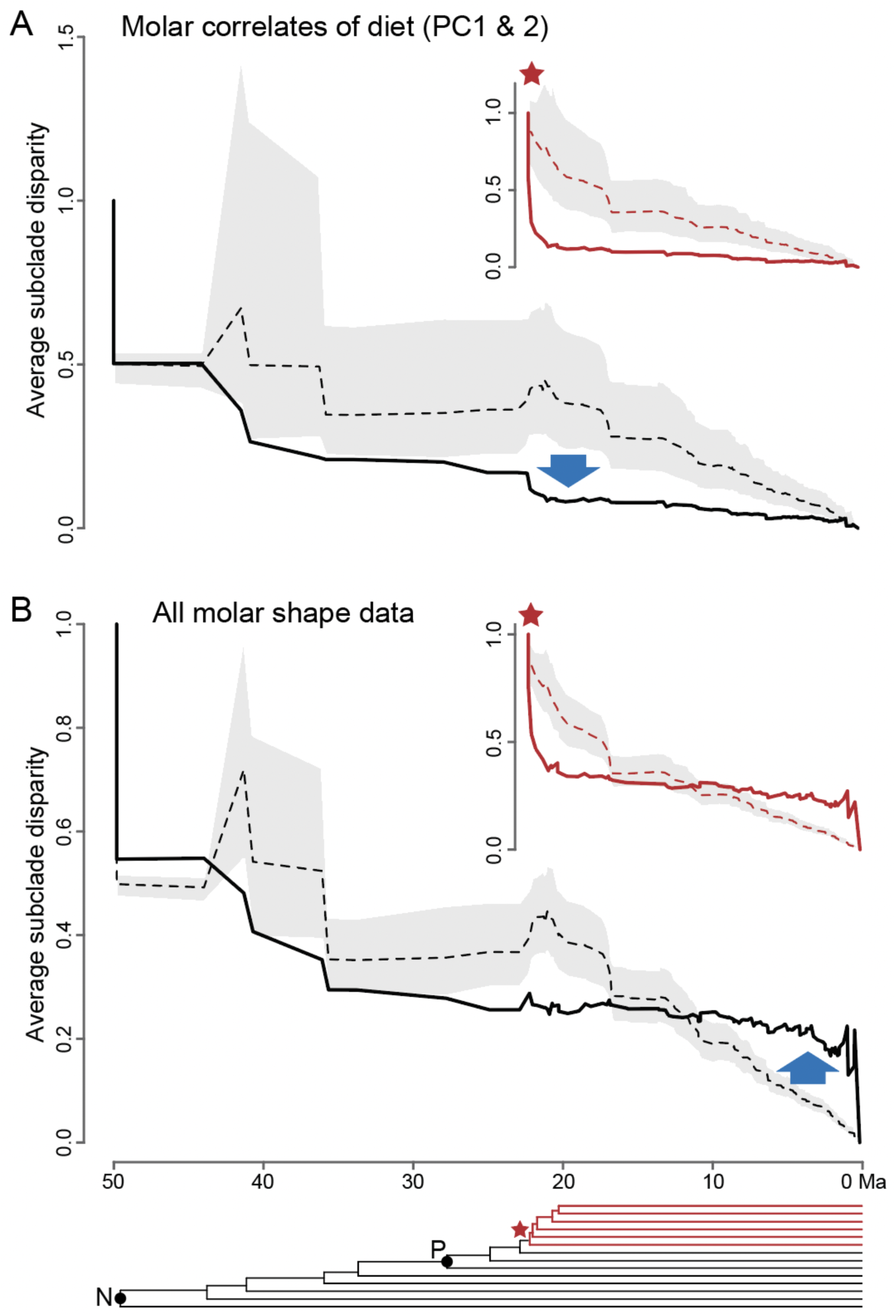
Subclade disparity-through-time plots when using only PC1–2 scores (diet correlates; *A*) and when using all PCs (*B*) from the geometric morphometrics analysis of molar shape. Solid lines are empirical data and dashed lines are simulated data with confidence intervals. The blue arrows highlight the major differences between the results of the two analyses. Analyses were performed using all Noctilionoidea (black) and only taxa defined by the node denoted by the red star (“Phyllostomidae*”, red), which includes most phyllostomids except for early branching faunivores (Fig. 1B). The phylogeny at the bottom is a simplified version of the phylogeny in Figure 1B, with Noctilionoidea (N) and Phyllostomidae (P) labeled.

**Table 1.**
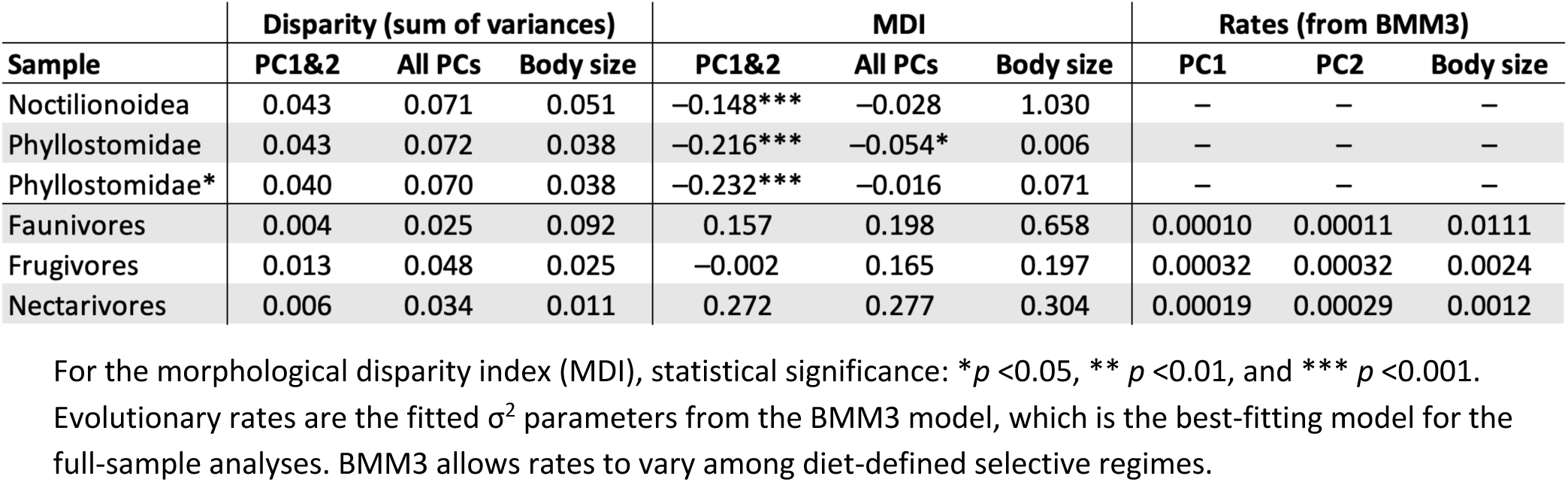
Morphological disparities and evolutionary rates of molar shape for focal clades and diet groups

| Sample | Disparity (sum of variances) |  |  | MDI |  |  | Rates (from BMM3) |  |  |
| --- | --- | --- | --- | --- | --- | --- | --- | --- | --- |
|  | PC1&2 | All PCs | Body size | PC1&2 | All PCs | Body size | PC1 | PC2 | Body size |
| Noctilionoidea | 0.043 | 0.071 | 0.051 | −0.148*** | −0.028 | 1.030 | — | — | — |
| Phyllostomidae | 0.043 | 0.072 | 0.038 | −0.216*** | −0.054* | 0.006 | — | — | — |
| Phyllostomidae* | 0.040 | 0.070 | 0.038 | −0.232*** | −0.016 | 0.071 | — | — | — |
| Faunivores | 0.004 | 0.025 | 0.092 | 0.157 | 0.198 | 0.658 | 0.00010 | 0.00011 | 0.0111 |
| Frugivores | 0.013 | 0.048 | 0.025 | −0.002 | 0.165 | 0.197 | 0.00032 | 0.00032 | 0.0024 |
| Nectarivores | 0.006 | 0.034 | 0.011 | 0.272 | 0.277 | 0.304 | 0.00019 | 0.00029 | 0.0012 |

For the morphological disparity index (MDI), statistical significance: \* $p < 0.05$ , \*\* $p < 0.01$ , and \*\*\* $p < 0.001$ . Evolutionary rates are the fitted $\sigma^2$ parameters from the BMM3 model, which is the best-fitting model for the full-sample analyses. BMM3 allows rates to vary among diet-defined selective regimes.

Node height tests indicate that morphological contrasts between nodes decrease with time in Phyllostomidae for PC1 (*p* = 0.009) and Phyllostomidae* for PC1 and PC2 (*p* = 0.034 and *p* = 0.027, respectively), which is consistent with an early burst. This pattern is not statistically significant when using the full Noctilionoid sample or any of the individual diet groups (Table S2).

Among diet groups, the levels of molar morphological disparity (variance of tip data) and stationary variance (calculated using parameters from the fitted OU1 model; see Methods) are greatest in frugivores and lowest in faunivores (Tables 1 and S2). Similarly, frugivores exhibit the fastest evolutionary rates (σ^2^) based on the fitted BMM3 model (Table 1). Basicranium width (i.e., body size) results exhibit a very different pattern, with faunivores having the greatest disparity, stationary variance, and evolutionary rates (Tables 1 and S2).

### Adaptive zone mapping

To help infer the adaptive landscape of noctilionoids, we performed two sets of model-fitting analyses to the molar shape data: (i) ‘full-sample analyses’ that examine the overall adaptive landscape of noctilionoids and (ii) ‘subgroup analyses’ that examine macroevolutionary patterns within diet groups and less inclusive clades (see Methods; Tables 2 and S3). For the full-sample analyses, we fitted single-regime, shift, and multiple-regime models to the entire sample of noctilionoids, with assigned selective regimes corresponding to diet categories. For analyses using molar data (either PC1–2 or PC1–6), the overwhelmingly best-fitting model is BMM3, a multiple-regime BM model that allows rates to vary between regimes and uses a three-diet classification scheme (faunivory, frugivory, and nectarivory). The second-best-fitting model is BMM4, which uses a four-diet classification scheme with omnivores as a separate regime/diet group. However, the considerably better fit of the BMM3 model suggests that there is not a distinct ecomorphotype associated with omnivory, which is consistent with morphological and performance studies of bat upper molars (Santana et al. 2011) and mammalian jaws (Grossnickle 2020). The best-fitting shift model for PC1–2 is OUEB, which models a shift from a constrained OU process in faunivores to an EB in frugivores and nectarivores. For the analyses of basicranium width (body size), OUBMi is the best-fitting model. OUBMi is the ‘release and radiate’ model of Slater (2013), which allows for a rate change with the shift between modes.

The presence of three adaptive zones is independently supported by the results of the *phyloEM* model-fitting analyses, which do not include *a priori* regime (diet) assignments. For the PC1–2 analysis, five regime shifts were identified, and these largely correspond to major diet shifts (Fig. S4). Two shifts are at the base of the nectarivore clades (Glossophaginae and Loncophyllinae), and three shifts are likely related to frugivory. There are no shifts within faunivore clades, supporting our inference that these taxa represent a single, ancestral adaptive zone. The *phyloEM* analysis using additional PCs (PC1–6) shows evidence for a greater number of regime shifts (>50), which are less correlated with diet compared to the shifts in the PC1–2 analysis.

For the subgroup analyses, we fitted single-regime models (BM1, OU1, EB) to subgroups within Noctilionoidea, including Phyllostomidae, Phyllostomidae*, and groups that only include individuals of one diet (faunivores, frugivores, and nectarivores). An EB model is the best-fitting model in analyses that (i) include multiple diet groups and (ii) use molar correlates of diet (PC1–2; Fig. 3A–C, Table S3). For the EB model fit to Phyllostomidae*, the rate decay half-life is 3.4 million years, which indicates that seven rate half-lives have elapsed during the clade’s history and suggests a relatively strong early burst pattern (Slater and Pennell 2014). When using additional molar data (PC1–6), the strong support for an EB model disappears, suggesting that evolutionary patterns of additional molar traits strongly conflict with an early burst pattern. The EB model performs poorly in all other analyses, except for frugivore molar correlates of diet, in which it is the second-best performing model with a ΔAICc of <2 (Table S3, Fig. 3E).

**Fig. 3.**
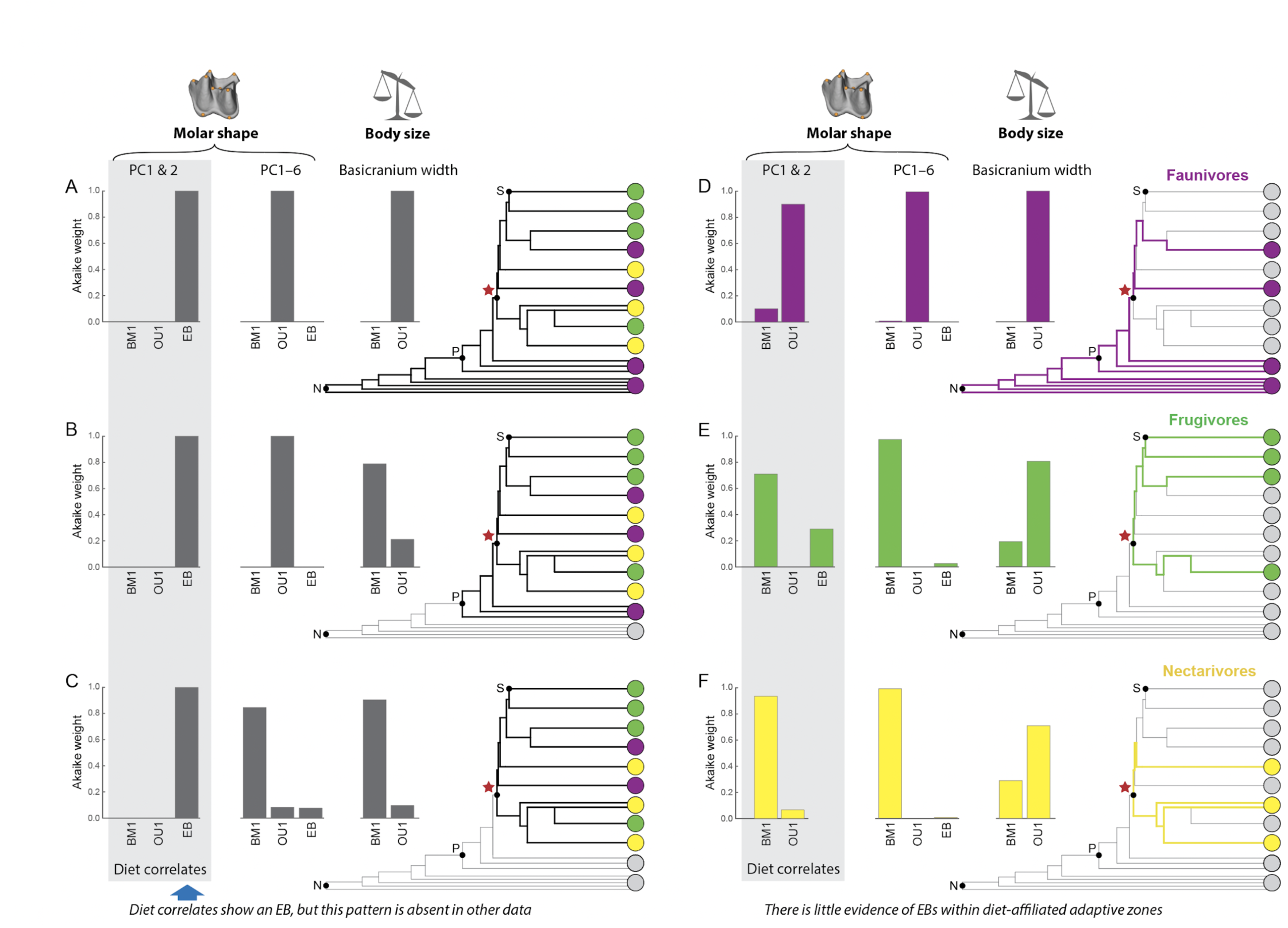
Evolutionary model-fitting analyses testing the influence of node choice (*A–C*) and examining patterns within diet-affiliated adaptive zones (*D–F*). These ‘subgroup analyses’ only include single-regime models. Prior to analyses, we pruned the phylogeny to the focal lineages for each analysis, and the excluded lineages are gray tree branches. We use the three-diet scheme (excluding omnivory as a separate group) because multiple-regime models using the three-diet scheme outperformed the four-diet-scheme models (Table 1). Each analysis was repeated using three morphological datasets (molar shape PC1–2, molar shape PC1–6, and log-transformed basicranium width). EB model results are excluded when the decay rate parameter is zero because the model collapses to a BM model. The phylogeny is a simplified version of that in Figure 1B; sister taxa with shared diets are condensed into single lineages and some lineages are illustrated as being paraphyletic for simplicity. Abbreviations: BM1, single-regime Brownian motion model; EB, early burst model; N, Noctilionoidea; OU1, single-regime Ornstein-Uhlenbeck model; P, Phyllostomidae; ‘red star’, Phyllostomidae*; S, Stenodermatinae.

For basicranium width, the EB model always collapses to a BM model (i.e., the rate decay parameter is zero), and OU1 is the best-fitting model for all three diet groups (Fig. 3).

## DISCUSSION

The observed macroevolutionary patterns of phyllostomid molars are consistent with predictions of the hierarchical adaptive radiation hypothesis (Osborn 1902, Simpson 1944, Simpson 1953, Slater and Friscia 2019). The phyllostomid radiation involved (i) a dramatic higher-level radiation of lineages into novel, diet-affiliated adaptive zones and (ii) lower-level radiations of lineages within adaptive zones. The tempos and modes of evolution of these two levels of radiation vary considerably, with the higher-level radiation involving relatively faster, more extreme morphological changes consistent with an early burst pattern, as predicted by Simpson (1944). We conceptualize our interpretation of this hierarchical radiation in Figure 4, and the following two subsections discuss evidence for these patterns.

**Fig. 4.**
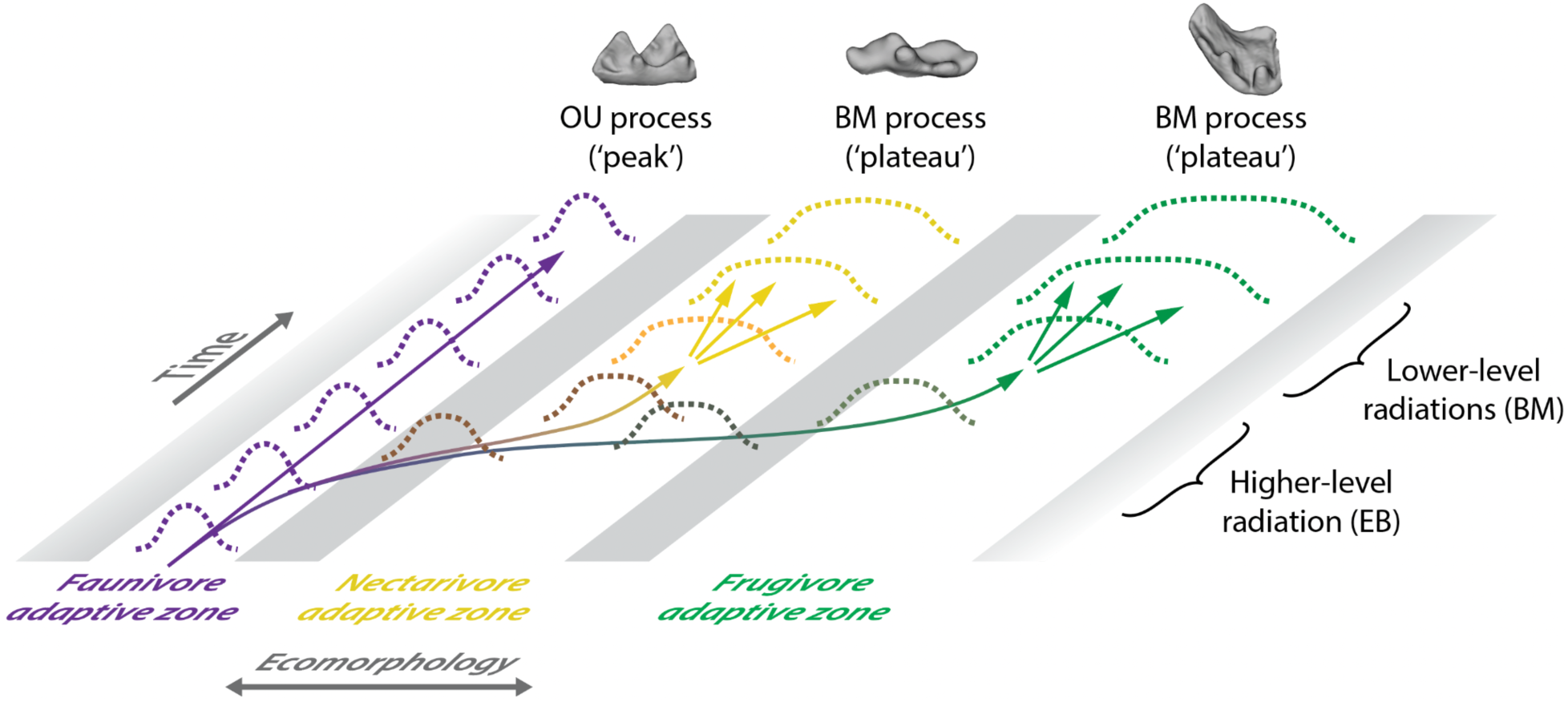
Conceptualization of the hierarchical radiation of phyllostomid bats, inspired by illustrations of ‘quantum evolution’ and adaptive zones in Simpson (1944). The higher-level radiation of phyllostomids exhibits an early burst (EB) pattern (Simpson’s ‘explosive phase’), but within-zone radiations are best described by Brownian motion (BM) or Ornstein-Uhlenbeck (OU) processes (Simpson’s ‘normal phase’).

### Higher-level radiation – early burst

The adaptive landscape of Noctilionoidea likely includes three (or more) broad, diet-affiliated adaptive zones (Fig. 4). This inference is supported by the separation of diet groups in morphospace (Fig. 1C), the especially strong fit of the BMM3 model to molar shape data (Table 2), the identification of regime shifts by the *phyloEM* analysis that largely correspond to major diet shifts near the Phyllostomidae* node (Fig. S4), and pANOVAs and pMANOVAs that show significant differences in molar shapes among diet groups (especially when using the three-diet scheme; Table S1). Further, the scarcity of molar shapes in intermediate regions of the morphospace between diet groups suggests the presence of adaptive valleys between zones (Fig. 1C). This initial radiation also likely included the evolution of sanguivores (Freeman 1998, Baker et al. 2012, Rossoni et al. 2017), which were excluded from analyses due to their extremely derived, reduced molars (Fig. S2). Thus, a fourth adaptive zone is likely present for sanguivores. Our results are consistent with inferences of the phyllostomid adaptive landscape based on jaw shape (Monteiro and Nogueira 2011, Arbour et al. 2019, Clavel and Morlon 2020). Monteiro and Nogueira (2011) also found evidence that vertivores and insectivores occupy separate adaptive zones in mandible morphology. We were unable to explicitly test this scenario due to a small sample size of vertebrate-eating taxa, but molar shapes of vertivores and insectivores do not appear to separate into different adaptive zones in our morphospace (Figs. 1C and S1).

**Table 2.**
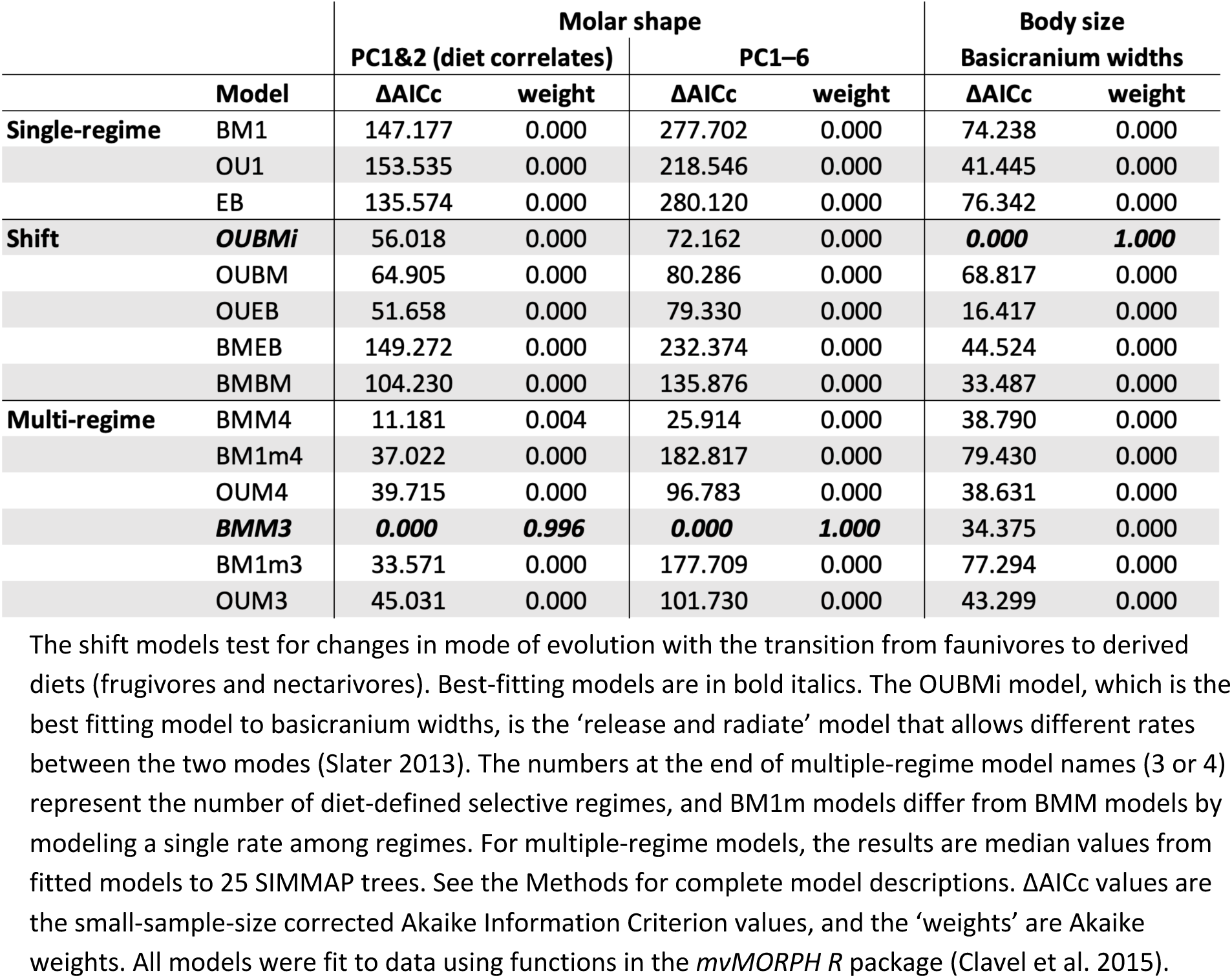
Evolutionary model-fitting analyses that test the overall adaptive landscape of Noctilionoidea (‘full-sample analyses’).

|  | Model | Molar shape |  |  |  | Body size |  |
| --- | --- | --- | --- | --- | --- | --- | --- |
|  |  | PC1&2 (diet correlates) |  | PC1–6 |  | Basicranium widths |  |
| | | $\Delta AICc$ | weight | $\Delta AICc$ | weight | $\Delta AICc$ | weight |
| Single-regime | BM1 | 147.177 | 0.000 | 277.702 | 0.000 | 74.238 | 0.000 |
|  | OU1 | 153.535 | 0.000 | 218.546 | 0.000 | 41.445 | 0.000 |
|  | EB | 135.574 | 0.000 | 280.120 | 0.000 | 76.342 | 0.000 |
| Shift | ***OUBMi*** | 56.018 | 0.000 | 72.162 | 0.000 | **0.000** | **1.000** |
|  | OUBM | 64.905 | 0.000 | 80.286 | 0.000 | 68.817 | 0.000 |
|  | OUEB | 51.658 | 0.000 | 79.330 | 0.000 | 16.417 | 0.000 |
|  | BMEB | 149.272 | 0.000 | 232.374 | 0.000 | 44.524 | 0.000 |
|  | BMBM | 104.230 | 0.000 | 135.876 | 0.000 | 33.487 | 0.000 |
| Multi-regime | BMM4 | 11.181 | 0.004 | 25.914 | 0.000 | 38.790 | 0.000 |
|  | BM1m4 | 37.022 | 0.000 | 182.817 | 0.000 | 79.430 | 0.000 |
|  | OUM4 | 39.715 | 0.000 | 96.783 | 0.000 | 38.631 | 0.000 |
|  | ***BMM3*** | **0.000** | **0.996** | **0.000** | **1.000** | 34.375 | 0.000 |
|  | BM1m3 | 33.571 | 0.000 | 177.709 | 0.000 | 77.294 | 0.000 |
|  | OUM3 | 45.031 | 0.000 | 101.730 | 0.000 | 43.299 | 0.000 |

The shift models test for changes in mode of evolution with the transition from faunivores to derived diets (frugivores and nectarivores). Best-fitting models are in bold italics. The OUBMi model, which is the best fitting model to basicranium widths, is the 'release and radiate' model that allows different rates between the two modes (Slater 2013). The numbers at the end of multiple-regime model names (3 or 4) represent the number of diet-defined selective regimes, and BM1m models differ from BMM models by modeling a single rate among regimes. For multiple-regime models, the results are median values from fitted models to 25 SIMMAP trees. See the Methods for complete model descriptions. $\Delta AICc$ values are the small-sample-size corrected Akaike Information Criterion values, and the 'weights' are Akaike weights. All models were fit to data using functions in the *mvMORPH* R package (Clavel et al. 2015).

Multiple lines of evidence suggest that the invasion of the frugivore and nectarivore adaptive zones occurred during a Simpsonian early burst (i.e., ‘explosive phase’) that involved very rapid ecomorphological change (Figs. 2 and 3, Tables 1 and S3). An early burst pattern at or near the Phyllostomidae node is evident in both the subclade disparity-through-time plots (Fig. 2) and the subgroup model-fitting analyses (Fig. 3). EB models best fitted the molar data for clades that include multiple diet groups (Fig. 3A–C), and the statistically significant MDI results (Table 1) and relatively short EB model rate decay half-lives (3.4 million years for Phyllostomidae*; Table S2) provide further evidence of a strong early burst. Although an EB model is best-fitting among single-regime models at all tested phylogenetic levels (Fig. 3A–C), results from additional analyses (node height test, disparity-through-time, and EB decay rate half-lives; Fig. 2, Tables 1 and S2) suggest that the EB specifically occurred near the Phyllostomidae or Phyllostomidae* node. For instance, when using only molar correlates of diet (PC1–2), a distinct decrease in subclade disparity is present at the base of Phyllostomidae* (Fig. 2A), which is where most dietary diversity begins to arise in Phyllostomidae (Fig. 1B). This suggests an early burst pattern in traits associated with diet at Phyllostomidae*, with subclade disparity rapidly decreasing as lineages shifted to new adaptive zones associated with frugivory and nectarivory.

Multiple-regime models outperform EB models in the ‘full sample’ analyses (Table 2), which is consistent with previous studies in which multiple-regime models better explain patterns of radiating clades – in agreement with a Simpsonian early burst (e.g., Mahler et al. 2013, Arbour et al. 2019). One potential reason for this result is that morphological convergence resulting from shifts between adaptive zones can weaken support for EB models (Slater and Pennell 2014) in favor of multiple-regime models (e.g., Mahler et al. 2013). Multiple phyllostomid lineages independently evolved both nectarivory and frugivory (Rojas et al. 2011; Fig. 1B), resulting in morphological convergence among these lineages (Fig. 1C). Nonetheless, the EB model is the best-fitting single-regime model and the OUEB model is the best-fitting shift model (Table 2, Fig. 3A–C), indicating that among the simpler classes of models there is strong support for an early burst.

In ‘subgroup analyses,’ the strong fits of EB models to molar correlates of diet could be due in part to relying on only two PCs, which could bias results toward an EB model (Uyeda et al. 2015). However, our primary conclusion (early burst in Phyllostomidae but not within individual adaptive zones) is supported by additional analyses, including the MDI and node height tests (Tables 1 and S2). Further, PC1–2 correspond closely with functional traits (e.g., relative cusp heights) that are linked to diet (e.g., Boyer et al. 2008, López-Aguirre et al. 2022, Villalobos-Chaves and Santana 2022), supporting our decision to use PC1–2 as functionally relevant traits. Finally, the model-fitting analyses are multivariate and incorporate correlation between PCs, which is less problematic than fitting univariate models to individual PCs (Clavel et al. 2015, Uyeda et al. 2015, Adams and Collyer 2018). Thus, we believe that our central conclusions are robust to potential biases associated with fitting models to PCs.

### Lower-level radiations – unique tempos and modes within subgroups

In contrast to the higher-level radiation, diversifications within diet-affiliated adaptive zones do not show strong evidence of early bursts (Fig. 3D–F). Instead, results of subgroup model-fitting analyses suggest that evolutionary patterns within frugivores and nectarivores are best described by a BM process (Fig. 3, Tables 2 and S3). Although the EB model is the second-best-fitting model in the frugivore analysis (Fig. 3E; and see Dumont et al. 2012), the EB model’s rate decay half-live is relatively long (10.8 million years when using PC1–6; Table S2) with only two half-lives having elapsed, suggesting a weak early burst pattern (Slater and Pennell 2014). In addition, frugivores’ MDI (Table 1) and node height test results (Table S2) are both not significant and therefore inconsistent with a strong early burst. Although not analyzed here, the sanguivores have reduced molars that appear to be morphologically similar among species, and thus it is unlikely that they experienced an early burst in morphology.

The adaptive landscape topography varies among diet groups, and these differences may reflect varying functions of molar ecomorphotypes. Faunivore molars, which represent the ancestral ecomorphotype (Fig. 1), are best fitted by an OU model, suggesting an adaptive peak with strong selective pressures maintaining the ancestral, tribosphenic (or tribosphenic-like dilambdodont) molar morphology that has remained relatively unchanged during chiropteran history (e.g., Freeman 2000, Jones et al. 2021). This is supported by the phylomorphospace and disparity results, which show that faunivores occupy a relatively small region of morphospace, as well as exhibit relatively slow evolutionary rates (Fig. 1C, Tables 1 and S2). The lack of disparity in this ecomorphotype is possibly due to strong selective pressures related to maintaining a functional tribosphenic molar. The upper and lower molars require precise occlusion of multiple shearing crests to function in the breakdown of insect cuticle and/or slicing through flesh (Evans and Sanson 2003, Polly et al. 2005), and small morphological changes could disrupt proper function to a greater degree than similar changes in frugivores and nectarivores. Further, the ancestral tribosphenic molar morphology likely evolved early in mammalian history (ca. 160 Ma; e.g., Luo et al. 2011), possibly resulting in well-established developmental constraints. Although faunivores do not show evidence of continued morphological diversification with time, they may have still experienced ecological diversification. For instance, extant faunivores within Noctilionoidea include taxa with specialized diets such as piscivory and vertivory (Dataset S1). Thus, the tribosphenic faunivore molar may represent many-to-one mapping of *function to form*, in which a single molar morphology is well suited for multiple faunivorous diets.

In contrast to faunivores, the strong fits of BM models in frugivore-only and nectarivore-only analyses suggest that these diet groups occupy relatively broad adaptive zones, which could be conceptualized as adaptive ‘plateaus.’ The interpretation of broad adaptive plateaus is further supported by relatively high disparity levels and fast evolutionary rates compared to those of faunivores (Table 1). After early lineages of these diet groups crossed adaptive valleys and reached adaptive plateaus, we posit that they continued to diversify morphologically rather than be constrained to a specific adaptive peak (Fig. 1C). This is reflected in results of analyses involving molar traits that are not strongly linked to diet. For instance, in the disparity-through-time results (Fig. 2), PC1–2 show a distinct decrease in subclade disparity at Phyllostomidae* (blue arrow in Figure 1B). But this pattern disappears entirely when using all molar shape data (Fig. 2B), and subclades show elevated disparity until the Present (blue arrow in Figure 2B). Therefore, whereas molar traits associated with diet show a diversification near the base of Phyllostomidae, additional molar traits continued to diversify within adaptive zones, maintaining the prolonged elevated disparity. The traits that continue to diversify may be associated with finer-scale resource partitioning as lineages saturated adaptive zones.

Within frugivores, the relatively greater diversity of molar shapes could reflect adaptations for consumption of specific fruit types. For instance, phyllostomid frugivores may specialize in soft, small-seeded fruits (e.g., Fleming and Heithaus 1986); hard, large-seeded fruits (e.g., Villalobos-Chaves et al. 2020); seed predation (e.g., Villalobos-Chaves et al. 2016); or may have generalized fruit diets. Some molar features (e.g., rim on the labial side of the molar, large trigon basin; Freeman 1988) are posited to relate to specialization on these narrower diets. In contrast, nectarivores rely less on molars for feeding (although they may still masticate some insects and plant products; e.g., Rojas et al. 2018), as they use their tongues for ingesting nectar. Thus, there may have been a relaxation of selective pressures to maintain specific molar morphologies, allowing greater diversification via stochastic processes such as drift, which is supported by the especially strong fit of a BM model (Fig. 3F). Further, nectarivores are morphologically distinguished from other groups by their especially long jaws and rostra (Fig. S2, Hedrick et al. 2021), and their relatively long and narrow molars could be a byproduct of the evolutionary lengthening and narrowing of the jaw (Sadier et al. 2022).

Beyond the evidence highlighted here, a hierarchical pattern during adaptive radiations is also supported by the hypothesis that radiations occur in stages, with each stage associated with a different resource axis (Streelman and Danley 2003). However, this idea is based primarily on radiations at relatively smaller phylogenetic scales without major shifts between adaptive zones (Streelman and Danley 2003, Gavrilets and Losos 2009, Colombo et al. 2015, Slater 2022). Thus, our study provides novel insight on evolutionary processes at broader scales, with a focus on a critical resource axis (i.e., diet) that likely drove the early burst pattern.

### Crossing adaptive valleys

There is a scarcity of taxa occupying morphospace regions between the three diet-affiliated adaptive zones (Fig. 1C), which we interpret as adaptive valleys. This is consistent with Simpson’s (1944) view that intermediate morphologies between adaptive zones are evolutionarily unstable and will rapidly evolve toward adaptive zones (i.e., ‘quantum evolution’). However, this raises the question as to how early phyllostomid lineages crossed this valley during their initial radiation. We build on previous research that suggests early phyllostomids were omnivorous (Freeman 2000, Baker et al. 2012, Hall et al. 2021) and that evolutionary shifts from omnivory to derived diets (frugivory, nectarivory, sanguivory) resulted in especially strong selective pressures that drove rapid evolutionary changes (Dávalos 2007, Rojas et al. 2011, Dumont et al. 2012, Rossoni et al. 2017, Rossoni et al. 2019).

As an ecological strategy, omnivory may have helped lineages ‘bridge’ the gap between adaptive zones because their molars may exhibit morphological tradeoffs between faunivory and herbivory (Freeman 2000, Polly 2020). The ancestral tribosphenic molar morphology is common among faunivores, but it also includes a mortar-like talonid basin that permits grinding or crushing of plant material. Thus, omnivores and insectivores often have very similar molar morphologies (e.g., Santana et al. 2011, Grossnickle and Newham 2016), which is supported in our results – most omnivores in our sample occupy the faunivore region of morphospace (Fig. 1C), and model-fitting results do not show evidence of a unique omnivore morphotype/adaptive zone (Table 2). Nonetheless, as plants become the prominent component of a lineage’s diet, the molars may adapt to additional grinding/crushing, resulting in a morphology more like that of frugivorous phyllostomids. This suggests that the intermediate morphology between faunivores and frugivores (near the middle of our morphospace) may be an optimal tradeoff morphology for taxa with plant-dominated omnivorous diets. Freeman (2000) specifically suggested that the molar morphology of *Carollia* species, which are primarily plant-dominated omnivores, could represent this intermediate tradeoff morphology – this is supported by their central positioning in our morphospace (Fig. S1). As ancestral omnivores fully shifted to derived diets such as frugivory and nectarivory, they were likely exposed to strong selective pressures that drove major morphological changes in molar and cranial morphology (Dávalos 2007, Rojas et al. 2011, Dumont et al. 2012, Rossoni et al. 2017, Rossoni et al. 2019). This is supported by the strong relationship between major axes of variation (PC1–2) and diet (Fig. 1C, Table S1), in addition to PC1–2 accounting for more variance than expected by chance (via the broken-stick model; Fig. S3).

The path through the adaptive valley may have also been facilitated by developmental biases, which can either constrain or facilitate the adaptive evolution of morphology (Goswami et al. 2014, Harjunmaa et al. 2012, Couzens et al. 2021). For instance, the novel acquisition of cusps during the evolution of mammals has been linked to variation of signaling and growth factors as well as phenotypic drift, which allowed tooth morphologies to escape constraints and cross adaptive valleys (Couzens et al. 2021). Similarly, the evolution of bat molars seems to have overcome well-established developmental constraints, such as the inhibitory cascade model, to produce a large range of derived morphologies (Sadier et al. 2022).

## Conclusions

In their discussion of macroevolutionary patterns, Slater and Friscia (2019) concluded, “The critical question facing comparative evolutionary biologists is, we argue, not whether early burst adaptive radiation occurs but, rather, at what phylogenetic levels and in which niche traits it is most common.” Here, we answer that question for phyllostomids, finding that they experienced a family-level early burst radiation that was associated with dietary diversification. Importantly, this early burst pattern is only observed when examining molar traits that are strongly linked to diet. Body size, which is not correlated with diet in our sample, and non-diet-correlated molar traits show no evidence of an early burst, highlighting that results of comparative analyses may be especially sensitive to researchers’ choice of input data. We advocate the use of morphological traits that are strongly and functionally linked to ecology within focal clades, and we echo recent arguments against the reliance on body size data in macroevolutionary comparative studies (Slater and Friscia 2019, Grossnickle 2020, Slater 2022).

After the early burst, subsequent within-adaptive-zone diversifications among derived dietary groups (frugivores and nectarivores) occupy broad adaptive zones (or adaptive ‘plateaus’) associated with plant-based diets. The unique tempos and modes of evolution of different diet groups, including evidence of a constraining OU process for faunivores with ancestral tribosphenic molars, are likely linked to varying levels of selective pressures and functional demands on different molar ecomorphotypes. Thus, we interpret the phyllostomid radiation to be hierarchical, with an early burst only present at a higher-level scale that captures the invasion of new adaptive zones, followed by finer-scale niche partitioning as lineages diversified within newly invaded, broad adaptive zones (Fig. 4). This lends strong support to the hierarchical adaptive radiation hypothesis (Osborn 1902, Simpson 1944, Slater and Friscia 2019), and suggests that a hierarchical pattern may be a common, yet underappreciated, pattern among adaptive radiations.

## MATERIALS & METHODS

### Sample

We collected micro-computed tomography (μCT) scans of 405 noctilionoid bat skulls. We excluded specimens with worn or fractured first lower molars, resulting in a final sample of 315 specimens, representing 124 species and 56 genera (Dataset S1). The specimens are primarily from the American Museum of Natural History (AMNH), Burke Museum of Natural History & Culture (UWBM), and the University of Michigan Museum of Zoology (UMMZ). Scanning was performed at the University of Washington using Bruker’s Skyscan 1172 and Skyscan 1173 μCT scanners. Further, our sample includes scans obtained from the MorphoSource online repository (www.morphosource.org); many of these scans are available due to a recent effort to digitize bat diversity (Shi et al. 2018). The μCT image stacks were imported into 3DSlicer using SlicerMorph ImageStacks (Rolfe et al. 2021) and then converted to mesh models using 3DSlicer.

Dietary classifications of species are based on multiple sources (Dataset S1), including Rojas et al. (2018) and Lopez-Aguirre et al. (2022). We used two different classifications schemes: a three-diet scheme and a four-diet scheme. The four-diet scheme includes faunivory (broadly including insectivory, carnivory, and piscivory), frugivory, nectarivory, and omnivory. The three-diet scheme excludes omnivory as a category, and omnivorous species are classified into the other three diet groups based on their preferred food items and/or their ancestral diets (Dataset S1). We excluded sanguivores because their first lower molar is too derived for collection of landmarks for geometric morphometrics. See the Supplementary Methods (SI Appendix) and Dataset S1 for more information on diet classifications.

We collected our primary morphometric data (see below) on the first lower molar (m1). We chose the m1 because it is present in all taxa and is easily identifiable, and it has especially strong functional importance in bats (e.g., López-Aguirre et al. 2022, Santana et al. 2022). For body size analyses, we measured basicranium widths as proxies for body size (Curtis et al. 2020), as other cranial size metrics such as skull length are heavily influenced by diet (e.g., nectarivores have disproportionately long rostra). These were measured using the digital mesh models of noctilionoid skulls and were then natural log-transformed prior to analyses. Because complete skulls were not available for all scanned specimens, our sample for basicranium width analyses (*n* = 121 species) excluded *Enchisthenes hartii*, *Lonchophylla handleyi*, and *Vampyressa melissa*.

### Geometric morphometrics and analyses of PCs

We quantified the shapes of m1s using geometric morphometrics. The ten 3D landmark positions are shown in Figure 1A and described in the Supporting Information. Many frugivores possess especially derived features, including the occasional loss of cusps or gain of novel cusps, making the homology among some cusps uncertain (e.g., Freeman 1998, Velazco 2005). Therefore, we repeated analyses using a second landmarking scheme with alternative positions for the paraconid and metaconid landmarks for 20 frugivorous species (Supplementary Methods, Dataset S1). The results for analyses using this alternative landmarking scheme are very similar to primary results (Tables S4, S5, S6), and thus this issue is unlikely to influence our conclusions.

We accounted for differences in size and orientation of molars by performing a generalized Procrustes superimposition on the landmark data. For species with more than one specimen, we calculated species means using the Procrustes values, and we used these means in subsequent analyses. We ordinated and reduced the dimensionality of the Procrustes aligned coordinates using a principal component analysis (PCA). The Procrustes analysis and PCA were performed using functions in the *Geometric Morphometrics for Mathematica* package (Polly 2022). We produced a PC1–2 phylomorphospace plot (Fig. 1C) using the *phylomorphospace* function in the *phytools* package (Revell 2012) for *R* (R Core Team 2020). The phylogeny (Fig. 1B) used for the phylomorphospace and subsequent comparative analyses is a maximum clade credibility tree of 500 trees from the posterior distribution of the ‘completed trees’ analysis of Upham et al. (2019).

To determine which PCs are most closely linked to diet, we performed phylogenetic analyses of variance (pANOVAs) by regressing the scores of individual PCs against discrete diet categories using phylogenetic generalized least squares (PGLSs), and then applying an ANOVA test. Analyses were performed via the *gls* function in the *nlme* library (Pinheiro et al. 2022) for *R*, with the phylogenetic signal incorporated into the regressions estimated using maximum likelihood (i.e., the ‘lambda’ model setting). Further, pMANOVA were performed to test for differences in molar shape among diet groups using the *mvgls* and *manova.gls* functions of *mvMORPH* with 9999 permutations (Clavel and Morlon 2020). We performed a pMANOVA using all shape data (all Procrustes residuals) and using only PC1–2, and we repeated them using different evolutionary models (see Supplementary Results and Table S1).

### Evolutionary model-fitting analyses

We fit a suite of evolutionary models to the morphological datasets using the *mvMORPH R* library (Clavel et al. 2015). These include three classes of evolutionary models: single-selective-regime (or uniform) models, ‘shift’ models, and multiple-selective-regime models. The single-regime models are BM1 (‘random walk’), OU1 (random walk with a selection parameter; Hanson 1997, Butler and King 2004), and EB (random walk with a rate decay parameter; Freckleton and Harvey 2006, Harmon et al. 2010). Model support was determined using small-sample-size-corrected Akaike Information Criterion (AICc) values.

The shift models allow for a shift in mode of evolution (e.g., OUBM models a shift from OU to BM) between two selective regimes. Our goal was to examine a shift in mode of evolution as lineages ‘escape’ from the ancestral faunivore ecomorphotype (Fig. 1C). Thus, for each shift model, we treated faunivores as the initial (ancestral) evolutionary mode and non-faunivores (frugivores and nectarivores) as the derived mode.

In multiple-regime model-fitting analyses, the selective regimes correspond to diet group classifications. The three-regime models (BM1m3, BMM3, OUM3) use the three-diet classification scheme, and four-regime models (BM1m4, BMM4, OUM4) use the four-diet classification scheme, which includes omnivory as a dietary category. BM1m models allow the phylogenetic means to vary among regimes (“smean” set to FALSE) but keep the evolutionary rate (σ^2^) constant, whereas BMM models allow differences in rates between regimes. The reported evolutionary rates (Table 1) are the σ^2^ (rate-step) parameter from the three-regime BMM model (BMM3), which was the best fitting model to the full sample. Multiple-regime OU models (OUM) allow trait optima to vary among regimes, but σ^2^ and α remain constant. We drop the root state influence (i.e., root = “stationary”) because we are using an ultrametric tree without fossil evidence for the root state (Clavel et al. 2015).

We performed two sets of model-fitting analyses: ‘full-sample analyses’ and ‘subgroup analyses.’ We repeated each set of analyses using three morphological datasets: molar correlates of diet (PC1–2, 60% of variance), overall molar shape (PC1–6, 83% of variance), and basicranium widths. For the molar analyses, the evolutionary models are multivariate. We limited the ‘overall molar shape’ analyses to the first six PCs due to computational constraints. The two sets of analyses are summarized here:

i. Full-sample analyses. Using all species in our sample (*n* = 124), we fitted single-regime, shift, and multiple-regime evolutionary models to morphological data. The inclusion of multiple-regime models allowed us to test for the presence of multiple diet-affiliated adaptive zones, and to test whether omnivores occupy a distinct adaptive zone. For multiple-regime models, the ancestral diet histories were reconstructed on the phylogeny using stochastic character maps (i.e., SIMMAPs; Bollback 2006), produced using the *make.simmap* function of the *phytools* library (Revell 2012) and an equal rates model. The reported results are means from using 25 SIMMAP trees. For shift models, we mapped the ancestral regime (faunivores) and derived regime (frugivores and nectarivores) onto the phylogeny using the *paintSubTree phytools* function, with diet-shift positions based on the summary of SIMMAP reconstructions (Fig. 1B).
ii. Subgroup analyses. Using separate analyses for different subclades and specific diet groups (see Results and Fig. 3), we fitted single-regime evolutionary models to morphological data. Prior to analyses, we pruned the phylogeny to the focal lineages for each analysis. Because of smaller sample sizes and the focus on within-adaptive zone/regime patterns, we only fitted single-regime models in these analyses.

As a supplemental model-fitting analysis to independently test for regime shifts that are not based on *a priori* regime (diet) assignments, we used the *phyloEM* function and *scalar OU* model in the *PhylogeneticEM R* package (Bastide et al. 2018). This function automatically detects regime shifts while accounting for correlations among traits. We tested 100 α values for three parallel computations and repeated analyses using PC1–2 and PC1–6.

### Morphological disparity

We calculated morphological disparity by summing variances of the data (either PCs 1 and 2 for analyses of diet correlates or all Procrustes residuals for analyses of overall molar shape) for each focal group. Further, we measured the stationary variances (expected variance when an OU process has reached a stationary state; σ^2^/2α) of PC1 and PC2 scores for each of the diet groups, using the fitted OU1 evolutionary models and *stationary* function from the *mvMORPH R* package, which follows Bartoszek et al. (2012) for calculating the multivariate normal stationary distribution.

In addition, we analyzed disparity patterns using the *disparity-through-time* (*dtt*) function from the *geiger R* library (Pennell et al. 2014), which plots the average subclade disparity through time (Fig. 2) and provides the Morphological Disparity Index (MDI) statistic (Harmon et al. 2003, Slater et al. 2010). We used 1000 BM-evolved simulations for calculating the null disparity patterns. MDI is the area between the DTT for the data and the median of the simulations, with *p*-values calculated as the probability that the MDI is significantly more negative than expected under a BM model (Harmon et al. 2003, Slater et al. 2010).

As a supplemental test of an early burst pattern, we performed node height tests (Freckleton and Harvey 2006, Slater and Pennell 2014) by fitting a linear model to independent contrasts versus node heights via the *nh.test* function in the *geiger R* library (Pennell et al. 2014). A significant, negative relationship reflects greater contrasts earlier in the phylogeny, providing evidence for an early burst.

## ACKNOWLEDGEMENTS

Nancy Simmons and Marisa Surovy facilitated access to specimens in the mammal collection at the AMNH, and Jeff Bradley assisted with specimens at the UWBM. We thank Jeff J. Shi, Cody W. Thompson, and Daniel L. Rabosky for providing access to specimen scans through MorphoSource.org, with funding for their research assisted by the National Science Foundation (NSF) DEB 1501307. Funding for this study was provided by the NSF (IOS 2017738 to S. E. S. and IOS 2017803 to K. E. S.). For feedback and assistance, we thank Lucas Weaver, P. David Polly, Jennifer Gardner, Adam P. Summers, Graham Slater, P. David Polly, Murat Maga, Sara Rohlfe, William H. Brightly, and members of the Sharlene Santana lab.

## SUPPORTING INFORMATION APPENDIX for

### SUPPLEMENTARY METHODS

#### Landmarks

For the geometric morphometrics analyses, we used ten 3D landmarks of the first lower molar (m1), which are illustrated in Figure 1A. Five of the landmarks (1–4, 7) are tips of molar cusps and one landmark is at a specific position on a molar crest (6). The other landmarks (5, 8–10) are Type III (Bookstein 1997) landmarks that help capture the general shape of the molar. Similar landmarks have been used in previous studies, including in 3D analyses of treeshrew molars (Selig et al. 2019; all but our Landmark 10 are included in their study) and 2D analyses of fossil molars (Wilson 2013, Grossnickle and Newham 2016), which showed that these landmarks help capture dietary differences between taxa.

More information on these landmarks are provided here:

1. Paraconid tip. See discussion below on alternative landmarking schemes for an explanation of the cusp position when the paraconid is absent.
2. Protoconid tip.
3. Metaconid tip. See discussion below on alternative landmarking schemes for an explanation of the cusp position when the metaconid is absent or there are multiple candidate metaconid cusps.
4. Entoconid tip.
5. Deepest (or central) point of the talonid basin. For most molars, we positioned the landmark at the deepest point of the talonid basin. However, in taxa with especially flat ‘basins’ (e.g., many frugivores), we used the center of the talonid basin region, approximately equidistant from surrounding cusps.
6. Cristid obliqua. We used the mesial-most point of the cristid obliqua, which is at the junction with the protoconid.
7. Hypoconid tip.
8. Labial-most point (in occlusal view) at the base of the talonid region/hypconid. This landmark was placed on the cingulum if present.
9. Labial-most point (in occlusal view) at the base of the trigonid region/protoconid. This landmark was placed on the cingulum if present.
10. Lingual-most point (in occlusal view) at the base of the metaconid, approximately at the midpoint of the length of the molar. This landmark was placed on the cingulum if present.

#### Alternative landmarking scheme for frugivores

For most taxa in our dataset, the ten landmark positions are easily identifiable. However, many frugivorous taxa possess especially derived molar features, including the occasional loss of cusps or gain of novel cusps, making the homology among some cusps uncertain (e.g., Freeman 1998, and see the contradictory scoring of phylogenetic morphological characters in studies such as Wetterer et al. 2000 and Velazco 2005). In some cases, we made subjective decisions for the locations of the paraconid and metaconid landmarks, and when there was a lack of candidate cusps (i.e., the paraconid and/or metaconid appeared to have been evolutionarily lost), we placed landmark points in the presumed homologous region. For instance, the paraconid tends to be located at the anterior margin of mammalian molars, and thus when the paraconid was absent we generally placed the corresponding Landmark 1 at the anterior-most point of the occlusal surface of the molar (see below). Similarly, many frugivorous taxa have especially large protoconids that may have ‘engulfed’ nearby, smaller cusps, meaning that the paraconid and/or metaconid could comprise part of the base of the protoconid cusp and not exhibit discernible features for placement of landmarks.

To help account for this uncertainty in landmark locations among frugivores, we repeated analyses using two landmarking schemes with alternative positions for the paraconid and metaconid landmarks (Landmarks 1 and 3) for some frugivores (*n* = 20 species; Dataset S1). For the first, or ‘outer,’ landmarking scheme, we placed Landmark 3 (metaconid) on a posteromedial cusp when multiple candidate cusps were present or on the medial edge of the molar when there were no candidate cusps. We placed Landmark 1 (paraconid) at the anterior-most point of the occlusal surface of the molar in cases in which this position was questionable. For the second, or ‘inner,’ landmarking scheme, Landmark 3 was placed on a more anterolateral candidate cusp when there were multiple candidate metaconid cusps. When the metaconid was absent, Landmark 3 was placed on the medial surface of the base of the protoconid, with the assumption that the large protoconid may have ‘engulfed’ the smaller metaconid. Similarly, when the paraconid was absent, Landmark 1 was placed at the anterior base of the protoconid, which was often in a very similar position as the ‘outer’ landmark scheme position.

#### Diet classifications

Dataset S1 includes sources for species’ diets (full references are in this file) and notes on some classification decisions. We primarily relied on Lopez-Aguirre et al. (2022), which builds on data in Rojas et al. (2011), Rojas et al. (2012), and Rojas et al. (2018), as well as compendiums (Nowak 1999, Wilman et al. 2014). We used two different classifications schemes: a three-diet scheme and a four-diet scheme. The four-diet scheme includes faunivory (broadly including insectivory, carnivory, and piscivory), frugivory, nectarivory, and omnivory. The three-diet scheme excludes omnivory as a category. We excluded sanguivores from analyzes because their first lower molar is too derived for collection of landmarks for geometric morphometrics.

We acknowledge that categorizing taxa into discrete dietary categories can be subjective, especially for omnivorous taxa, and alternative diet assignments could influence results of analyses that require diet classifications (e.g., multiple-regime model-fitting analyses). However, we strived to circumvent this issue by repeating multiple-regime modeling analyses without *a prior* regime (diet) assignments (Fig. S4), and many of our central conclusions are supported by clade-wide analyses that do not differentiate diet groups (e.g., the disparity-through-time analyses; Fig. 3).

Most mammals, including noctilionoids, are omnivorous to some degree, consuming both animals and plants (Nowak 1999, Pineda-Munoz et al. 2014, Rojas et al. 2018, Reuter et al. 2023, Pellón et al. 2023). This makes the classification of omnivores especially challenging and subjective (Pineda-Munoz et al. 2014, Grossnickle 2020, Reuter et al. 2023). We chose to only classify species as omnivores if the animal and plant components of their diets are approximately equal. This is a more restrictive definition of omnivory than that used in Rojas et al. (2018). For the three-diet scheme, which excludes omnivory as a category, the omnivorous species are classified into the other three diet groups based on their preferred food items and/or their ancestral diets (see notes in Dataset S1).

Ideally, we would separate faunivores into multiple diet categories, such as vertivory and insectivory. However, the sample of vertivores is too small (only two or three species in our sample consume primarily vertebrates; Dataset S1, Rojas et al. 2018) to be appropriate for defining vertivory as a separate selective regime (e.g., Cooper et al. 2016).

Two species that are especially challenging to classify are *Phyllonycteris aphylla* and *Phyllostomus discolor*. *Phyllonycteris aphylla* is a very restricted and critically endangered species, and little is known about its diet. Rojas et al. (2018) classify it as a frugivore, but this is based on a single observation of individuals feeding on fruit (Genoways et al. 2005). Other members of the genus are nectarivores and its ancestral diet is nectarivory (Fig. 1B), so presumably its preferred diet is nectar that is supplemented with fruits and insects (e.g., Nowak 1999). Thus, we classify *Phyllonycteris aphylla* as a nectarivore for both the three-and four-diet schemes. For *Phyllostomus discolor*, we classify it as an omnivore for the four-diet scheme and a faunivore for the three-diet scheme. Data in Rojas et al. (2018) indicates that it is more herbivorous than insectivorous, and Quinche et al. (2022) highlights some soft tissue specializations for nectarivory. However, its skull and body size are not characteristic of specialized nectarivory, and it does include a substantial proportion of fruit and insects in its diet, with lots of seasonal and geographic variation (Fleming et al 1972, Nowak 1999, Kwiecinski 2006). Thus, we chose to classify it based on its ancestral (and likely dominant) diet of insectivory/faunivory (Fig. 1B, Fleming et al 1972).

Based on these considerations, *Phyllonycteris aphylla* and *Phyllostomus discolor* could be classified as a frugivore and nectarivore, respectively. Interestingly, the phylomorphospace trajectories for *Phyllonycteris aphylla* and *Phyllostomus discolor* indicate that they are evolving toward the morphospace regions of these alternative diets (Figs. 1B and S1). *Phyllonycteris aphylla* is the nectarivore that is closest to frugivores, and *Phyllostomus discolor* is much closer in morphospace to nectarivores compared to additional *Phyllostomus* species (Fig. S1). Thus, these lineages appear to be morphological intermediates between diet ecomorphotypes (and they may be crossing adaptive valleys; see Discussion). If we were use the alternative diet categories for these species in our comparative analyses, we do not expect that it would have a large impact on results.

## SUPPLEMENTARY RESULTS

**Fig. S1.**
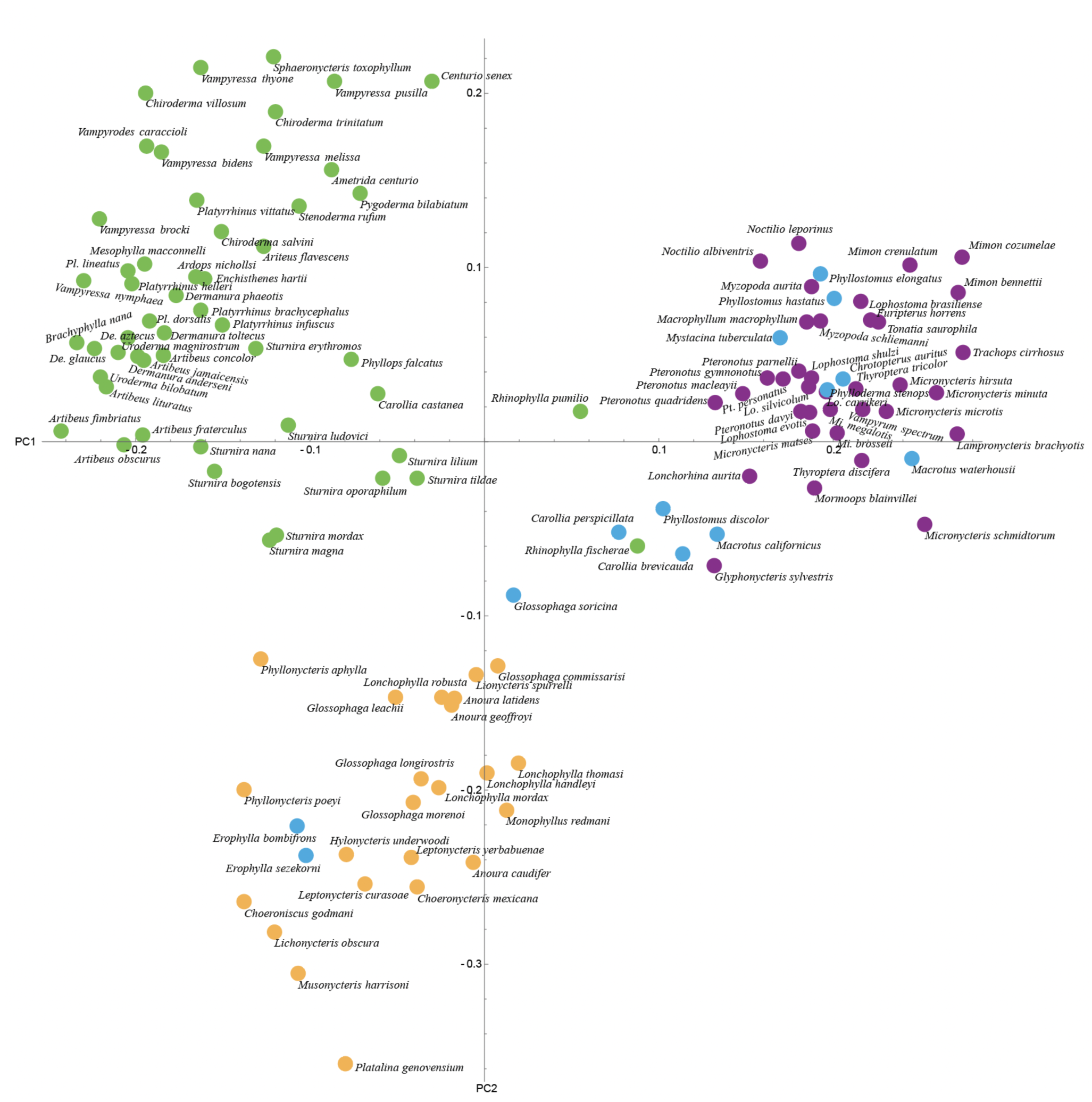
The PC1 and PC2 morphospace plot with species labels. This is the same plot as in Figure 1C, but with labels and without an added phylogeny.

**Fig. S2.**
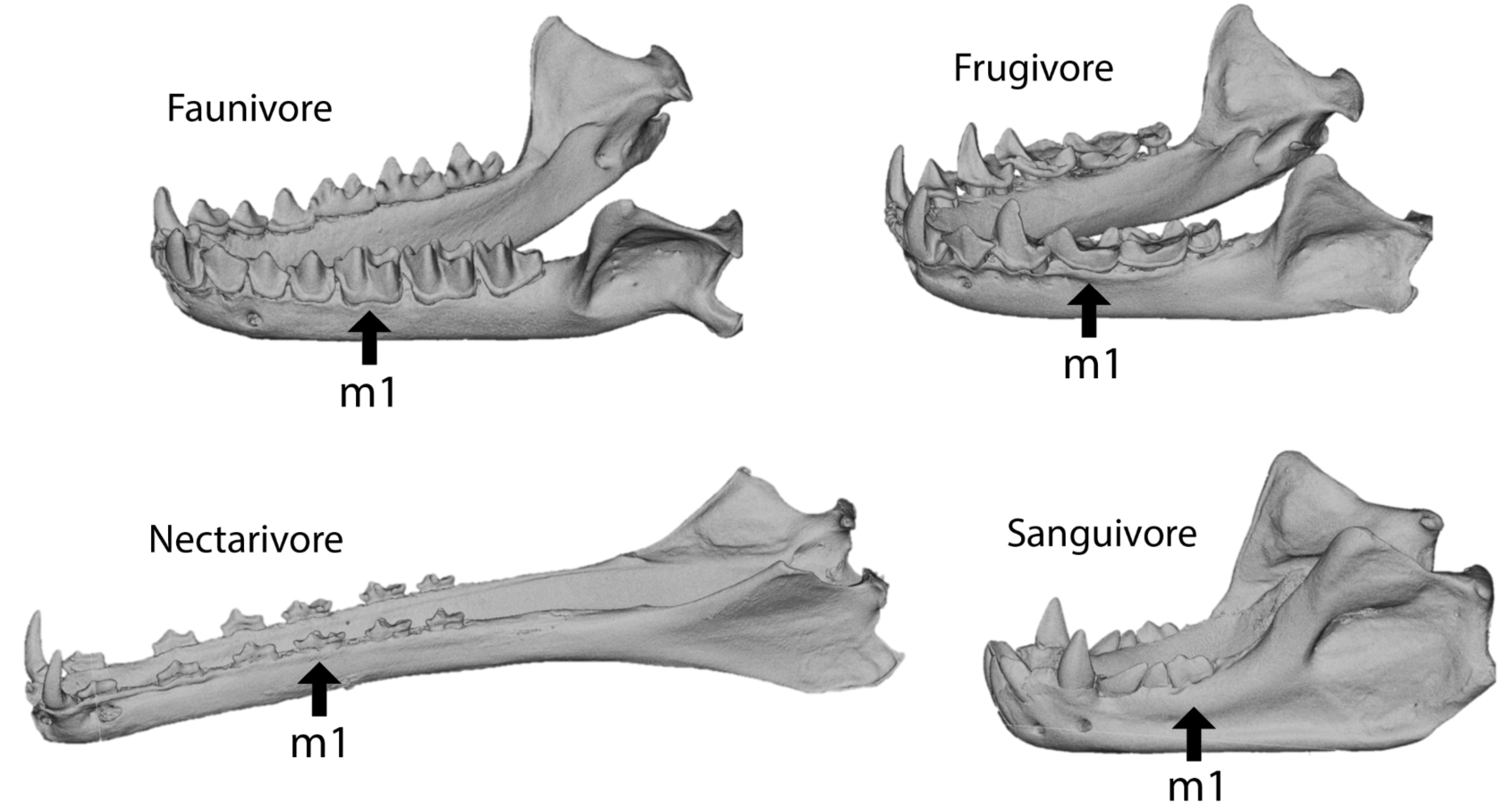
Representative jaws of the major diet groups examined in this study. Sanguivores were excluded due to their derived molars, which are relatively reduced and lack many of the cusps used as landmarks in the geometric morphometrics analyses of molar shape. The jaws are *Lophostoma silvicolum* (faunivore), *Platyrrhinus dorsalis* (frugivore), *Musonycteris harrisoni* (nectarivore), and *Diaemus youngi* (sanguivore).

**Fig. S3.**
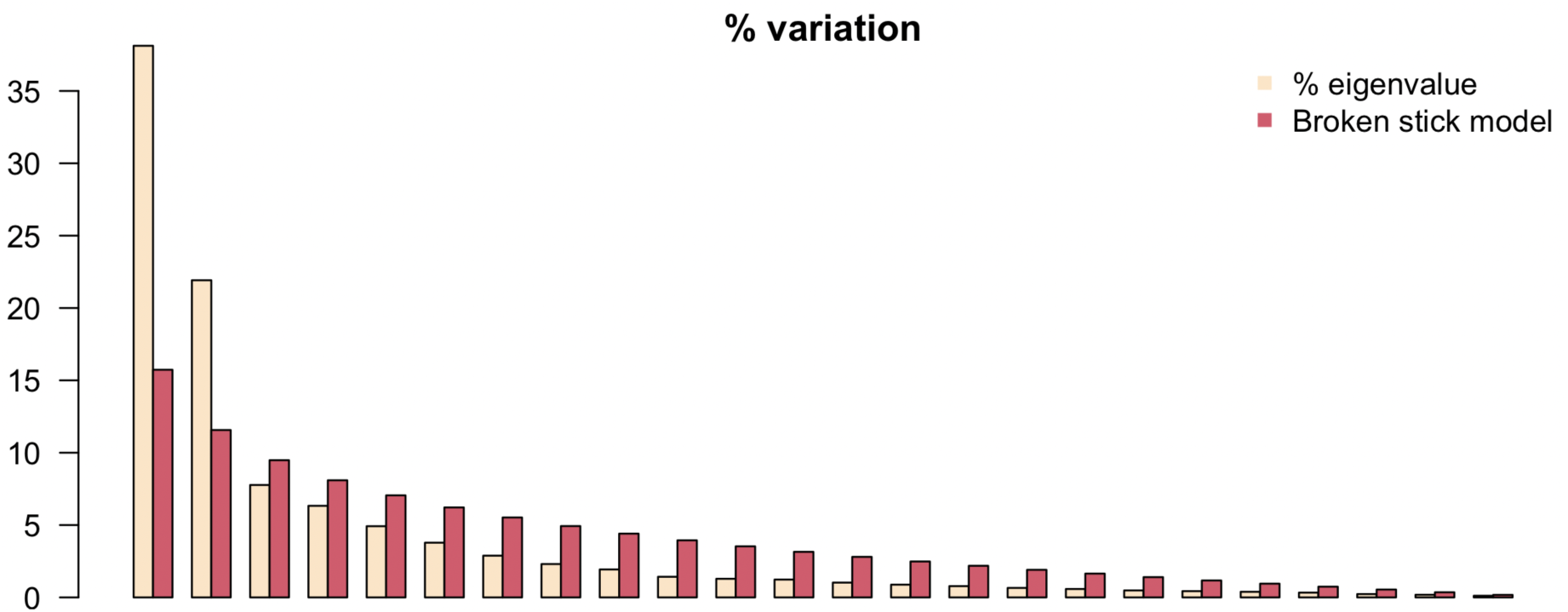
Broken-stick model for testing which principal components (PCs) explain a greater percentage of variance than is expected by chance. Note that the eigenvalues for the first two PCs are considerably greater than the variances of the broken stick model, whereas all other PCs have values that are less than the broken stick model.

**Table S1.**
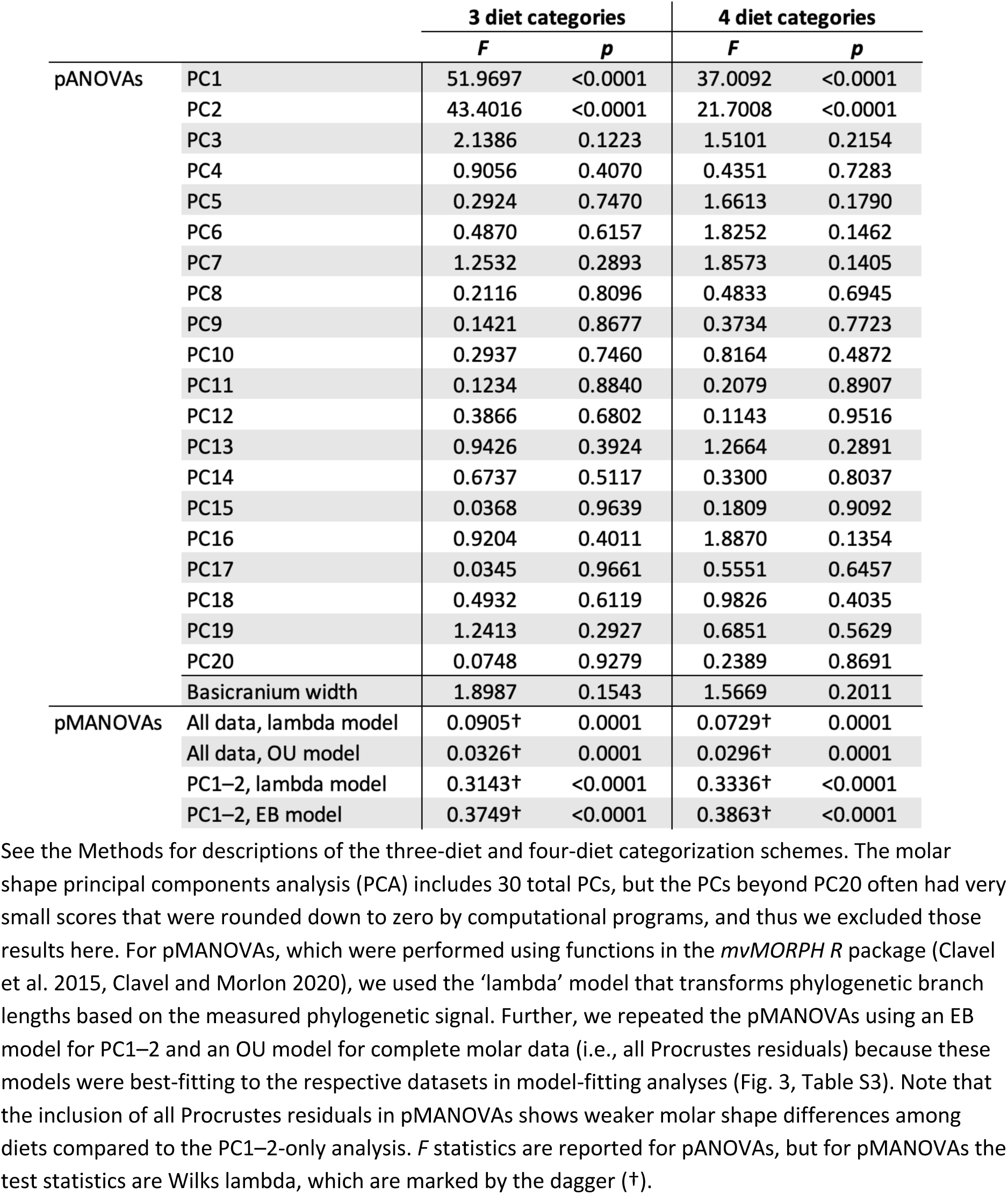
Summary statistics for phylogenetic one-way analyses of variance (pANOVAs) and phylogenetic multivariate analyses of variance (pMANOVAs).

**Table S2.**
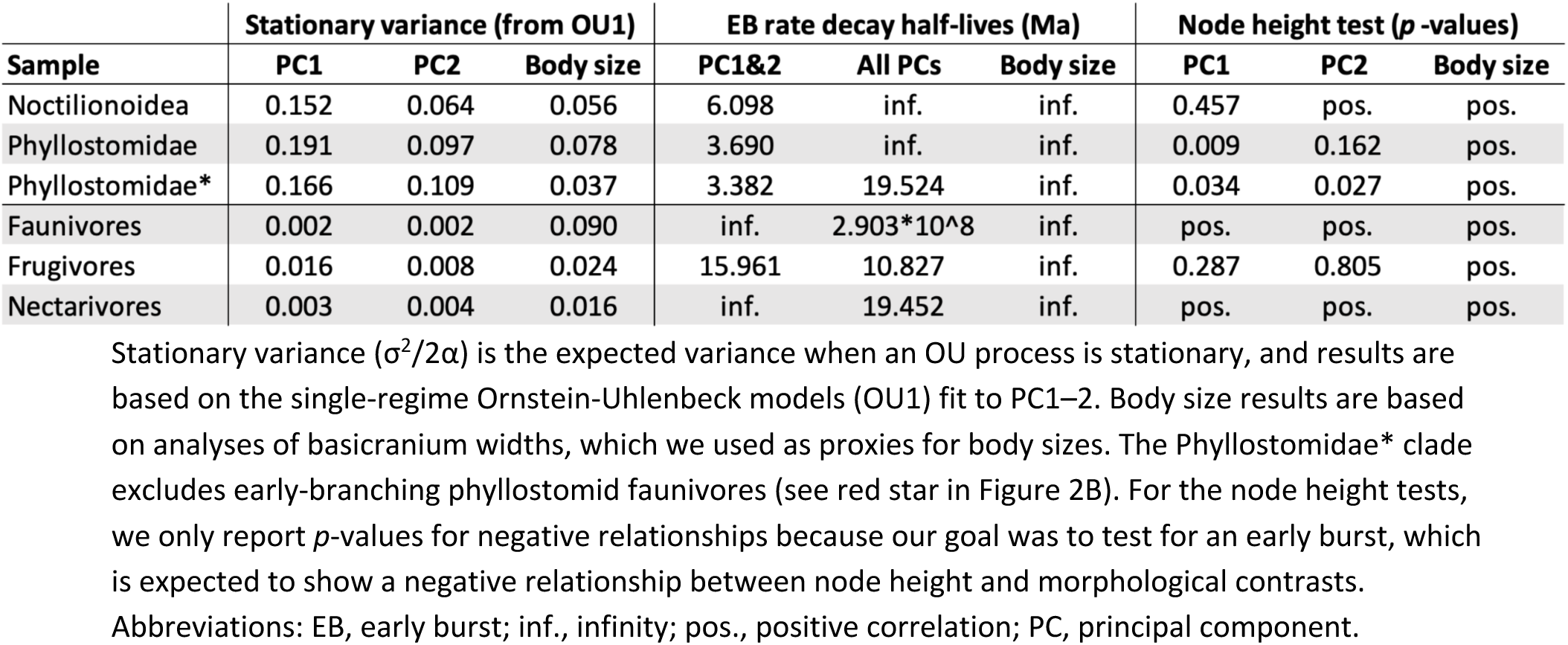
Results for supplemental measures of disparity (stationary variance) and tests of early burst (EB) patterns.

**Table S3.**
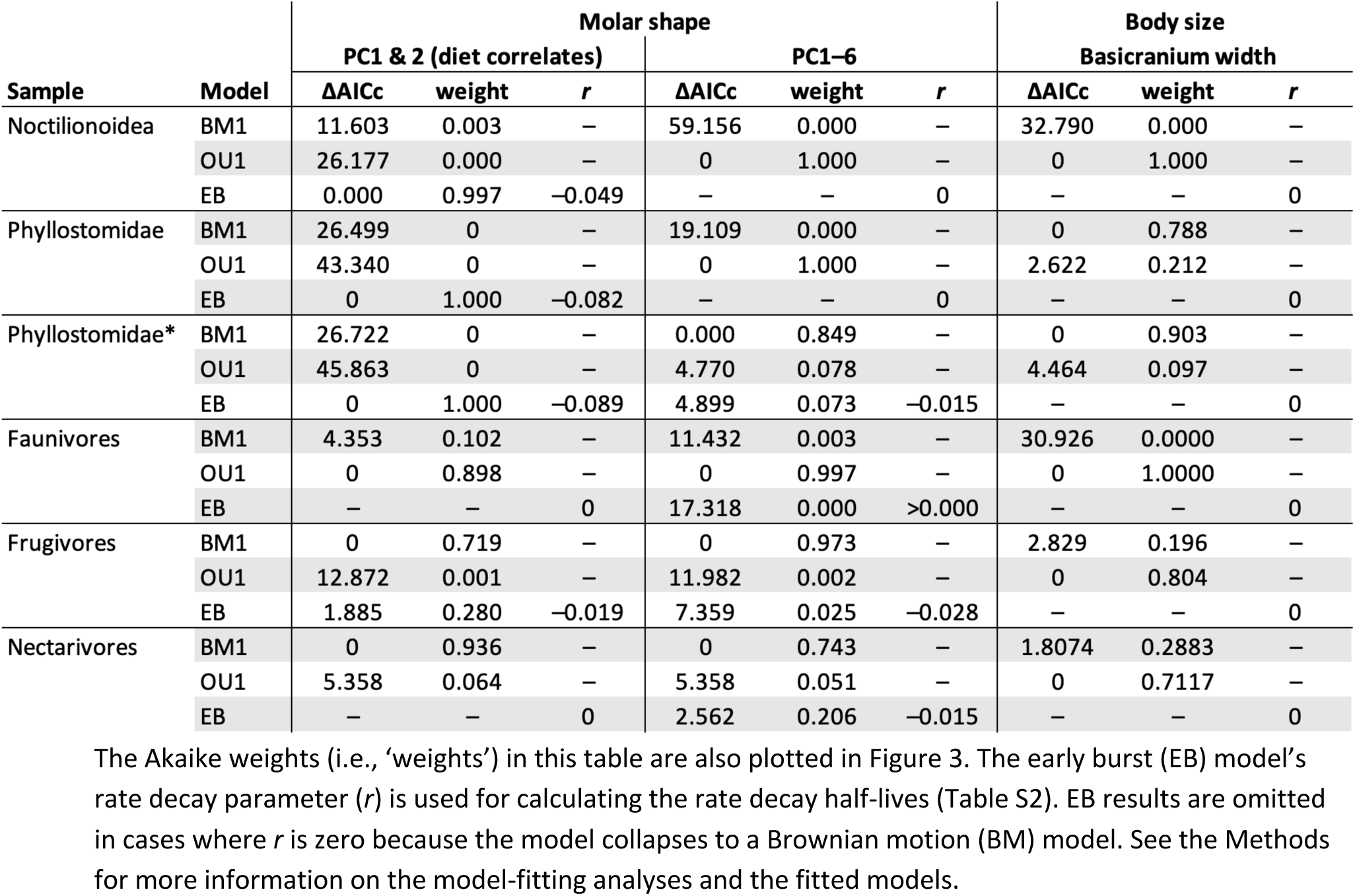
Evolutionary model-fitting results for the ‘subgroup analyses.’

**Table S4.**
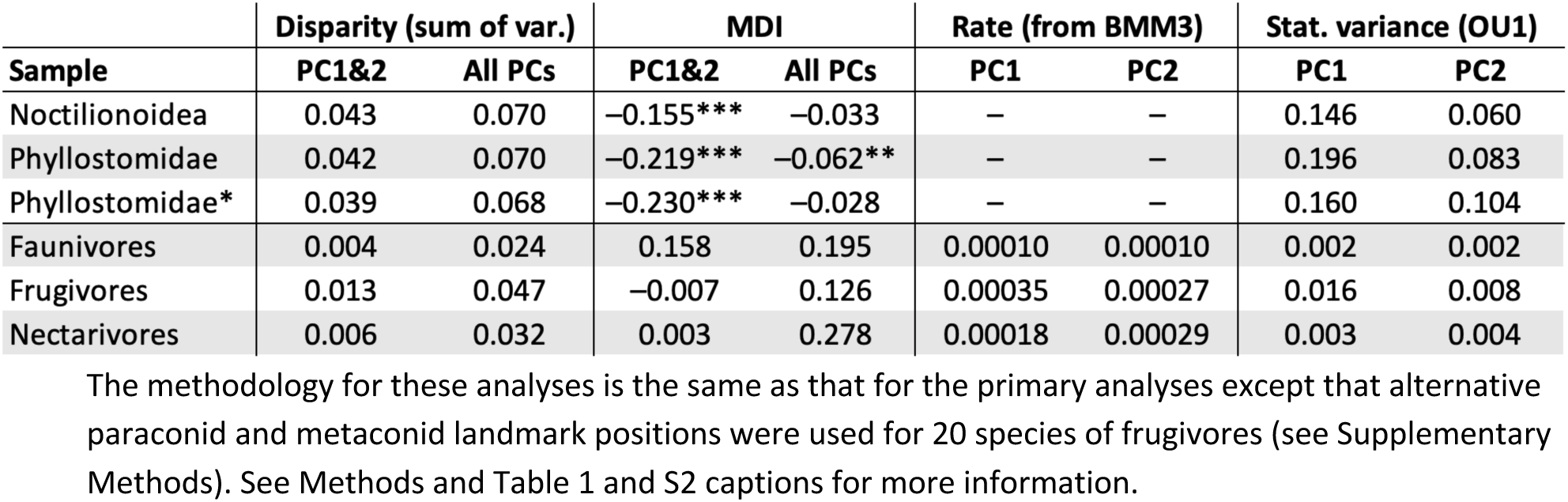
Disparity and evolutionary rate analyses using the alternative landmarking scheme for some frugivores.

**Table S5.**
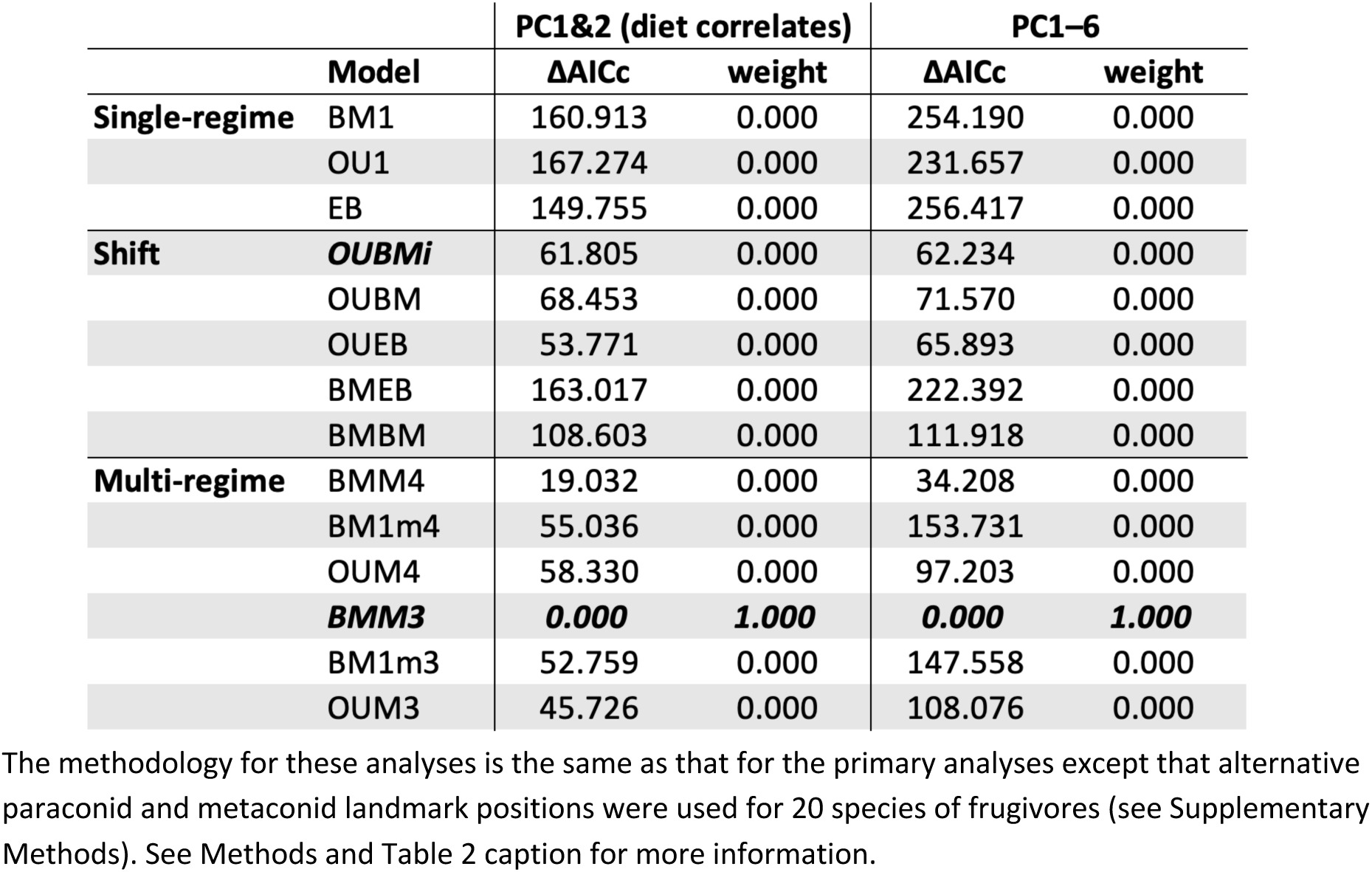
Evolutionary model-fitting results for the ‘full sample’ analyses using the alternative landmarking scheme.

**Table S6.**
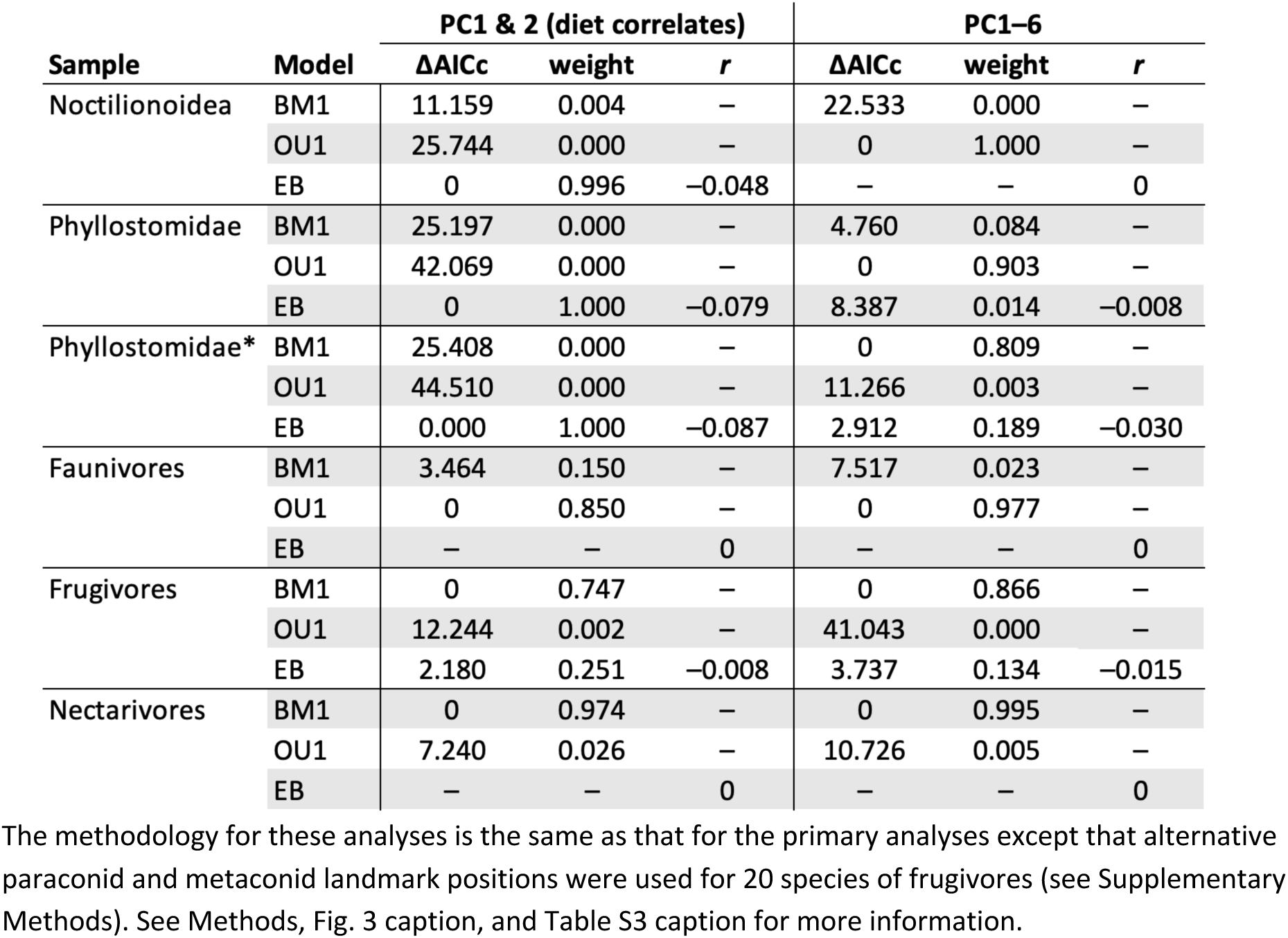
Evolutionary model-fitting results for the ‘subgroup’ analyses using the alternative landmarking scheme.

**Fig. S4.**
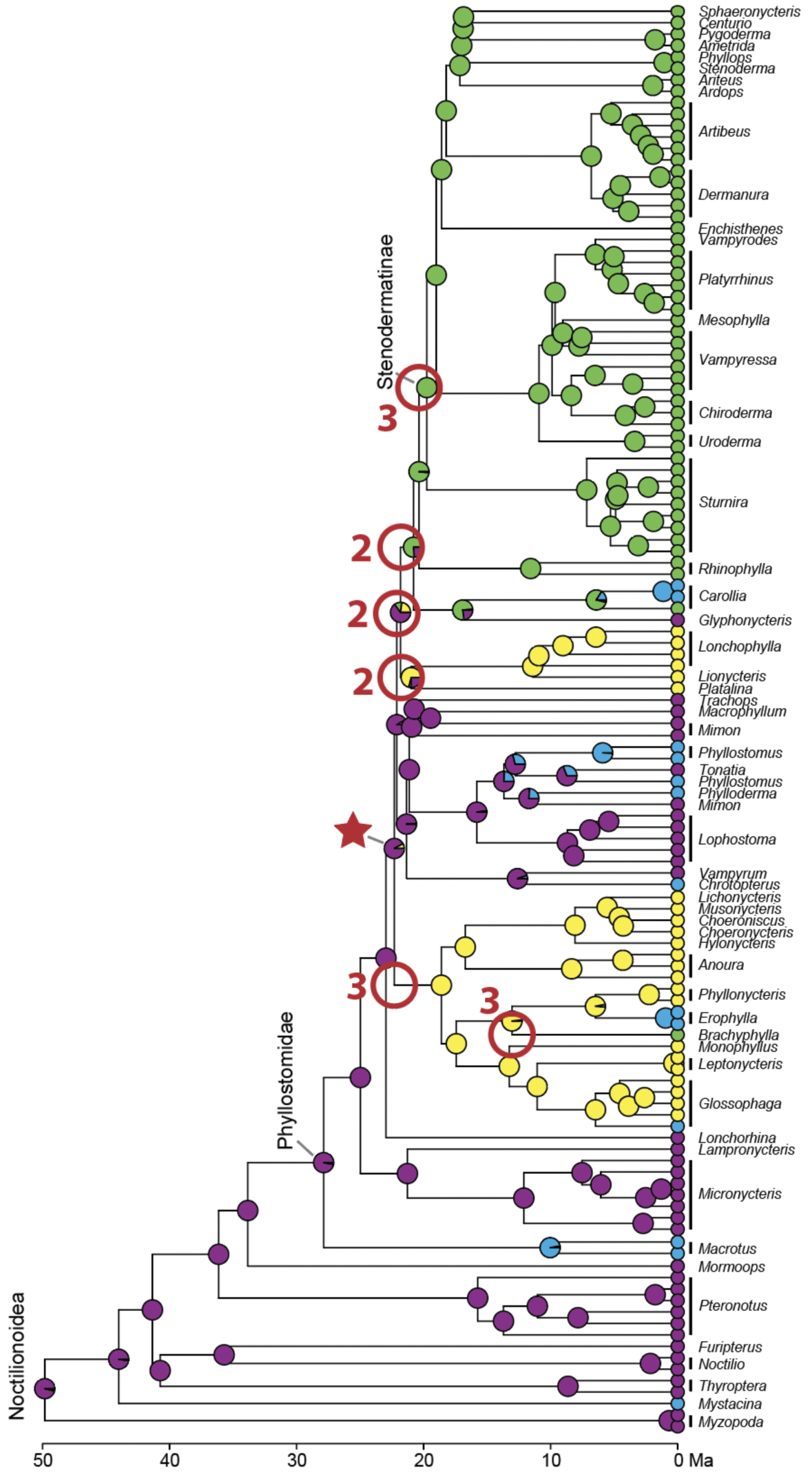
Model-fitting analyses for PC1–2 without *a priori* regime assignments via functions in the *PhylogeneticEM R* package (Bastide et al. 2018). The analysis found evidence of five regime shifts, but there were three equivalent solutions for shift positions, which included six positions in total (marked by red circles). The numbers next to the red circles indicate how many (out of three) equivalent solutions included shifts at those positions. Note that all shifts are within the Phyllostomidae* node and mostly correspond to dietary shifts, consistent with results for the ‘full sample’ model-fitting analyses with *a priori* regime assignments (Table 2). The *phyloEM* analysis results using PC1–6 indicated 11 regime shifts, but we are not illustrating them here because there were approximately 150 equivalent solutions with various shift locations throughout the tree.

## REFERENCES

1. Adams, D. C., Collyer, M. L. (2018). Multivariate phylogenetic comparative methods: evaluations, comparisons, and recommendations. Systematic Biology, 67, 14–31.

2. Arbour, J. H., Curtis, A. A., Santana, S. E. (2019). Signatures of echolocation and dietary ecology in the adaptive evolution of skull shape in bats. Nature Communications, 10, 2036.

3. Arbour, J. H., López-Fernández, H. (2013). Ecological variation in South American geophagine cichlids arose during an early burst of adaptive morphological and functional evolution. Proceedings of the Royal Society B: Biological Sciences, 280, 20130849.

4. Baker, R. J., O. R. P. Bininda-Emonds, H. Mantilla-Meluk, C. A. Porter, R. A. van den Bussche. 2012. Molecular time scale of diversification of feeding strategy and morphology in New World Leaf-Nosed Bats (Phyllostomidae): a phylogenetic perspective. Pp. 385–409 in G. F.Gunnell, ed. Evolutionary history of bats: fossils, molecules and morphology. Cambridge Univ. Press, Cambridge, U.K.

5. Bartoszek K., Pienaar J., Mostad P., Andersson S., Hansen T.F. 2012. A phylogenetic comparative method for studying multivariate adaptation. Journal of Theoretical Biology, 314, 204–215.

6. Bastide, P., Ané, C., Robin, S., & Mariadassou, M. (2018). Inference of adaptive shifts for multivariate correlated traits. Systematic Biology, 67, 662–680.

7. Bollback, J. P. (2006). SIMMAP: stochastic character mapping of discrete traits on phylogenies. BMC Bioinformatics, 7, 1–7.

8. Boyer, D. M. (2008). Relief index of second mandibular molars is a correlate of diet among prosimian primates and other euarchontan mammals. Journal of Human Evolution, 55, 1118–1137.

9. Burnham, K. P., & Anderson, D. R. (2004). Multimodel inference: understanding AIC and BIC in model selection. Sociological methods & research, 33, 261–304.

10. Butler, M. A., & King, A. A. (2004). Phylogenetic comparative analysis: a modeling approach for adaptive evolution. The American Naturalist, 164, 683–695.

11. Clavel, J., Escarguel, G., & Merceron, G. (2015). mvMORPH: an R package for fitting multivariate evolutionary models to morphometric data. Methods in Ecology and Evolution, 6, 1311–1319.

12. Clavel, J., Morlon, H. (2020). Reliable phylogenetic regressions for multivariate comparative data: illustration with the MANOVA and application to the effect of diet on mandible morphology in phyllostomid bats. Systematic biology, 69, 927–943.

13. Colombo, M., Damerau, M., Hanel, R., Salzburger, W., & Matschiner, M. (2015). Diversity and disparity through time in the adaptive radiation of Antarctic notothenioid fishes. Journal of Evolutionary Biology, 28, 376–394.

14. Couzens, A. M., Sears, K. E., & Rücklin, M. (2021). Developmental influence on evolutionary rates and the origin of placental mammal tooth complexity. Proceedings of the National Academy of Sciences, 118, e2019294118.

15. Curtis, A. A., Smith, T. D., Bhatnagar, K. P., Brown, A. M., & Simmons, N. B. (2020). Maxilloturbinal aids in nasophonation in horseshoe bats (Chiroptera: Rhinolophidae). The Anatomical Record, 303(1), 110–128.

16. Dávalos, L. M. (2007). Short-faced bats (Phyllostomidae: Stenodermatina): A Caribbean radiation of strict frugivores. Journal of Biogeography, 34, 364–375.

17. Dumont, E. R., Davalos, L. M., Goldberg, A., Santana, S. E., Rex, K., Voigt, C. C. (2012). Morphological innovation, diversification and invasion of a new adaptive zone. Proceedings of the Royal Society B: Biological Sciences, 279, 1797–1805.

18. Dumont, E. R., Samadevam, K., Grosse, I., Warsi, O. M., Baird, B., & Davalos, L. M. (2014). Selection for mechanical advantage underlies multiple cranial optima in new world leaf-nosed bats. Evolution, 68, 1436–1449.

19. Erwin, D. H. (1992). A preliminary classification of evolutionary radiations. Historical Biology, 6, 133–147.

20. Erwin, D. H. (2007). Disparity: morphological pattern and developmental context. Palaentology, 50, 57– 73.

21. Erwin, D. H. (2015). Novelty and innovation in the history of life. Current Biology, 25, R930–R940.

22. Evans, A. R., & Sanson, G. D. (2003). The tooth of perfection: functional and spatial constraints on mammalian tooth shape. Biological Journal of the Linnean Society, 78, 173–191.

23. Feilich, K. L., López-Fernández, H. (2019). When does form reflect function? Acknowledging and supporting ecomorphological assumptions. Integrative and Comparative Biology, 59, 358–370.

24. Fleming, T. H., Dávalos, L. M., & Mello, M. A. (Eds.). (2020). Phyllostomid bats: a unique mammalian radiation. University of Chicago Press.

25. Fleming, T.H. & Heithaus, E.R. (1986) Seasonal Foraging Behavior of the Frugivorous Bat Carollia perspicillata. Journal of Mammalogy, 67, 660–671.

26. Foote, M. (1994). Morphological disparity in Ordovician-Devonian crinoids and the early saturation of morphological space. Paleobiology, 20, 320–344.

27. Freeman, P.W. (1988). Frugivorous and animalivorous bats (Microchiroptera) – dental and cranial adaptations. Biological Journal of the Linnean Society, 33, 249–272.

28. Freeman, P. W. (1998). Form, function, and evolution in skulls and teeth of bats; in: Bat Biology and Conservation, T. H. Kunz, Ed. Smithsonian Institution Press, Washington, DC, 140–156.

29. Freeman, P. W. (2000). Macroevolution in Microchiroptera: recoupling morphology and ecology with phylogeny. Mammalogy Papers: University of Nebraska State Museum, 8.

30. Freckleton, R. P., Harvey, P. H. (2006). Detecting non-Brownian trait evolution in adaptive radiations. PLoS Biology, 4, e373.

31. Gavrilets, S., Losos, J. B. (2009). Adaptive radiation: contrasting theory with data. Science, 323, 732–737.

32. Givnish, T. J. (2015). Adaptive radiation versus ‘radiation’ and ‘explosive diversification’: why conceptual distinctions are fundamental to understanding evolution. New Phytologist, 207, 297–303.

33. Goswami, A., Smaers, J. B., Soligo, C., & Polly, P. D. (2014). The macroevolutionary consequences of phenotypic integration: from development to deep time. Philosophical Transactions of the Royal Society B: Biological Sciences, 369, 20130254.

34. Grossnickle, D. M. (2020). Feeding ecology has a stronger evolutionary influence on functional morphology than on body mass in mammals. Evolution, 74, 610–628.

35. Grossnickle, D. M., Newham, E. (2016). Therian mammals experience an ecomorphological radiation during the Late Cretaceous and selective extinction at the K–Pg boundary. Proceedings of the Royal Society B: Biological Sciences, 283, 20160256.

36. Grossnickle, D. M., Smith, S. M., Wilson, G. P. (2019). Untangling the multiple ecological radiations of early mammals. Trends in Ecology & Evolution, 34, 936–949.

37. Gunnell, G. F., Simmons, N. B., & Seiffert, E. R. (2014). New Myzopodidae (Chiroptera) from the Late Paleogene of Egypt: emended family diagnosis and biogeographic origins of Noctilionoidea. PLoS One, 9, e86712.

38. Harjunmaa, E., Kallonen, A., Voutilainen, M., Hämäläinen, K., Mikkola, M. L., & Jernvall, J. (2012). On the difficulty of increasing dental complexity. Nature, 483, 324–327.

39. Hall, R. P., et al. (2021). Find the food first: an omnivorous sensory morphotype predates biomechanical specialization for plant based diets in phyllostomid bats. Evolution, 75, 2791–2801.

40. Halliday, T. J. D., Goswami, A. (2016). Eutherian morphological disparity across the end-Cretaceous mass extinction. Biological Journal of the Linnean Society, 118, 152–168.

41. Hansen, T. F. (1997). Stabilizing selection and the comparative analysis of adaptation. Evolution, 51, 1341–1351.

42. Harmon, L. J., et al. (2010). Early bursts of body size and shape evolution are rare in comparative data. Evolution, 64, 2385–2396.

43. Harmon, L. J., Schulte, J. A., Larson, A., Losos, J. B. (2003). Tempo and mode of evolutionary radiation in iguanian lizards. Science, 301, 961–964.

44. Hedrick, B. P., Dumont, E. R. (2018). Putting the leaf-nosed bats in context: a geometric morphometric analysis of three of the largest families of bats. Journal of Mammalogy, 99, 1042–1054.

45. Hedrick, B. P., et al. (2020). Morphological diversification under high integration in a hyper diverse mammal clade. Journal of Mammalian Evolution, 27, 563–575.

46. Hughes, M., Gerber, S., Wills, M. A. (2013). Clades reach highest morphological disparity early in their evolution. Proceedings of the National Academy of Sciences, 110, 13875–13879.

47. Jackson, D. A. (1993). Stopping rules in principal components analysis: a comparison of heuristical and statistical approaches. Ecology, 74, 2204–2214.

48. Jones, M. F., Li, Q., Ni, X., & Beard, K. C. (2021). The earliest Asian bats (Mammalia: Chiroptera) address major gaps in bat evolution. Biology Letters, 17, 20210185.

49. López-Aguirre, C., Hand, S. J., Simmons, N. B., Silcox, M. T. (2022). Untangling the ecological signal in the dental morphology in the bat superfamily Noctilionoidea. Journal of Mammalian Evolution, 29, 531–545.

50. Losos, J. B. (2011). Lizards in an evolutionary tree: ecology and adaptive radiation of anoles. University of California Press. Berkeley, CA.

51. Luo, Z. X., Yuan, C. X., Meng, Q. J., & Ji, Q. (2011). A Jurassic eutherian mammal and divergence of marsupials and placentals. Nature, 476, 442–445.

52. Mahler, D. L., Ingram, T., Revell, L. J., & Losos, J. B. (2013). Exceptional convergence on the macroevolutionary landscape in island lizard radiations. Science, 341, 292–295.

53. Mitchell, J. S. (2015). Extant-only comparative methods fail to recover the disparity preserved in the bird fossil record. Evolution, 69, 2414–2424.

54. Monteiro, L. R., & Nogueira, M. R. (2010). Adaptive radiations, ecological specialization, and the evolutionary integration of complex morphological structures. Evolution, 64, 724–744.

55. Monteiro, L. R., & Nogueira, M. R. (2011). Evolutionary patterns and processes in the radiation of phyllostomid bats. BMC Evolutionary Biology, 11, 1–23.

56. Osborn, H. F. (1902). The law of adaptive radiation. The American Naturalist, 36, 353–363.

57. Pennell, M. W., et al. (2014). geiger v2.0: an expanded suite of methods for fitting macroevolutionary models to phylogenetic trees. Bioinformatics, 30, 2216–2218.

58. Pincheira-Donoso, D., Harvey, L. P., Ruta, M. (2015). What defines an adaptive radiation? Macroevolutionary diversification dynamics of an exceptionally species-rich continental lizard radiation. BMC Evolutionary Biology, 15, 1–13.

59. Pinheiro, J. C., Bates, D. M., R Core Team (2023). nlme: linear and nonlinear mixed effects models. R package version 3.1–162.

60. Polly, P. D. (2022). Geometric morphometrics for Mathematica, version 12.4. Department of Geological Sciences, Indiana University.

61. Polly, P. D. (2020). Functional tradeoffs carry phenotypes across the valley of the shadow of death. Integrative and Comparative Biology, 60, 1268–1282.

62. Polly, P. D., Le Comber, S. C., & Burland, T. M. (2005). On the occlusal fit of tribosphenic molars: Are we underestimating species diversity in the Mesozoic? Journal of Mammalian Evolution, 12, 283–299.

63. Price, S. L., Etienne, R. S., Powell, S. (2016). Tightly congruent bursts of lineage and phenotypic diversification identified in a continental ant radiation. Evolution, 70, 903–912.

64. R Development Core Team (2020). R: A language and environment for statistical computing, version 4.1.3. R Foundation for Statistical Computing, Vienna, Austria.

65. Revell, L. J. (2012). phytools: an R package for phylogenetic comparative biology (and other things). Methods in Ecology and Evolution, 2, 217–223.

66. Rojas, D., Vale, A., Ferrero, V., & Navarro, L. (2011). When did plants become important to leaf-nosed bats? Diversification of feeding habits in the family Phyllostomidae. Molecular Ecology, 20, 22172228.

67. Rojas, D., Ramos Pereira, M. J., Fonseca, C., & Dávalos, L. M. (2018). Eating down the food chain: generalism is not an evolutionary dead end for herbivores. Ecology Letters, 21, 402–410.

68. Rolfe, S., et al. (2021). SlicerMorph: An open and extensible platform to retrieve, visualize and analyse 3D morphology. Methods in Ecology and Evolution, 12, 1816–1825.

69. Rossoni, D. M., Assis, A. P. A., Giannini, N. P., & Marroig, G. (2017). Intense natural selection preceded the invasion of new adaptive zones during the radiation of New World leaf-nosed bats. Scientific Reports, 7, 11076.

70. Rossoni, D. M., Costa, B. M., Giannini, N. P., & Marroig, G. (2019). A multiple peak adaptive landscape based on feeding strategies and roosting ecology shaped the evolution of cranial covariance structure and morphological differentiation in phyllostomid bats. Evolution, 73, 961–981.

71. Ruta, M., Wagner, P. J., Coates, M. I. (2006). Evolutionary patterns in early tetrapods. I. Rapid initial diversification followed by decrease in rates of character change. Proceedings of the Royal Society B: Biological Sciences, 273, 2107–2111.

72. Sadier, A., et al. (2018). Multifactorial processes underlie parallel opsin loss in neotropical bats. Elife, 7, e37412.

73. Sadier, A., et al. (2022). Bat teeth illuminate the diversification of mammalian tooth classes. bioRxiv, 2022–11. doi:10.1101/2021.12.05.471324.

74. Santana, S. E., Grosse, I. R., & Dumont, E. R. (2012). Dietary hardness, loading behavior, and the evolution of skull form in bats. Evolution, 66, 2587–2598.

75. Santana S. E., Dumont E. R. and Davis J. L. 2010. Mechanics of bite force production and its relationship to diet in bats. Functional Ecology, 24, 776–784.

76. Santana, S. E., Strait, S., Dumont, E. R. (2011). The better to eat you with: functional correlates of tooth structure in bats. Functional Ecology, 25, 839–847.

77. Santana, S. E., Grossnickle, D. M., Sadier, A., Patterson, E., & Sears, K. E. (2022). Bat Dentitions: A Model System for Studies at the Interface of Development, Biomechanics, and Evolution. Integrative and Comparative Biology, 62, 762–773.

78. Schluter, D. (2000). The ecology of adaptive radiation. OUP Oxford.

79. Shi, J. J., Westeen, E. P., Rabosky, D. L. (2018). Digitizing extant bat diversity: An open-access repository of 3D μCT-scanned skulls for research and education. PLoS One, 13, e0203022.

80. Shi, J. J., Westeen, E. P., & Rabosky, D. L. (2021). A test for rate-coupling of trophic and cranial evolutionary dynamics in New World bats. Evolution, 75, 861–875.

81. Simpson, G. G. (1944). Tempo and mode in evolution. Columbia University Press. New York.

82. Simpson, G. G. (1953). The major features of evolution. Columbia University Press. New York.

83. Slater, G. J. (2022). Topographically distinct adaptive landscapes for teeth, skeletons, and size explain the adaptive radiation of Carnivora (Mammalia). Evolution, 76, 2049–2066.

84. Slater, G. J. (2013). Phylogenetic evidence for a shift in the mode of mammalian body size evolution at the Cretaceous-Palaeogene boundary. Methods in Ecology and Evolution, 4, 734–744.

85. Slater, G. J., Friscia, A. R. (2019). Hierarchy in adaptive radiation: a case study using the Carnivora (Mammalia). Evolution, 73, 524–539.

86. Slater, G. J., Harmon, L. J., Alfaro, M. E. (2012). Integrating fossils with molecular phylogenies improves inference of trait evolution. Evolution, 66, 3931–3944.

87. Slater, G. J., Pennell, M. W. (2014). Robust regression and posterior predictive simulation increase power to detect early bursts of trait evolution. Systematic Biology, 63, 293–308.

88. Slater, G. J., Price, S. A., Santini, F., Alfaro, M. E. (2010). Diversity versus disparity and the radiation of modern cetaceans. Proceedings of the Royal Society B: Biological Sciences, 277, 3097–3104.

89. Stanchak, K. E., Arbour, J. H., Santana, S. E. (2019). Anatomical diversification of a skeletal novelty in bat feet. Evolution, 73, 1591–1603.

90. Streelman, J. T., & Danley, P. D. (2003). The stages of vertebrate evolutionary radiation. Trends in Ecology & Evolution, 18, 126–131.

91. Stroud, J. T., & Losos, J. B. (2016). Ecological opportunity and adaptive radiation. Annual Review of Ecology, Evolution, and Systematics, 47, 507–532.

92. Uyeda, J. C., Caetano, D. S., & Pennell, M. W. (2015). Comparative analysis of principal components can be misleading. Systematic Biology, 64, 677–689.

93. Velazco, P. M. (2005). Morphological phylogeny of the bat genus Platyrrhinus Saussure, 1860 (Chiroptera: Phyllostomidae) with the description of four new species. Fieldiana Zoology, 2005, 1–53.

94. Villalobos-Chaves, D., Melo, F.P.L. & Rodríguez-Herrera, B. (2020) Dispersal patterns of large-seeded plants and the foraging behaviour of a frugivorous bat. Journal of Tropical Ecology, 36, 94–100.

95. Villalobos-Chaves, D., Padilla-Alvárez, S. & Rodríguez-Herrera, B. (2016) Seed predation by the wrinkle-faced bat Centurio senex: A new case of this unusual feeding strategy in Chiroptera. Journal of Mammalogy, 97, 726–733.

96. Villalobos-Chaves, D., & Santana, S. E. (2022). Craniodental traits predict feeding performance and dietary hardness in a community of Neotropical free-tailed bats (Chiroptera: Molossidae). Functional Ecology, 36, 1690–1699.

97. Weaver, L. N., & Grossnickle, D. M. (2020). Functional diversity of small-mammal postcrania is linked to both substrate preference and body size. Current Zoology, 66, 539–553.

98. Wilson, G. P., Evans, A. R., Corfe, I. J., Smits, P. D., Fortelius, M., Jernvall, J. (2012). Adaptive radiation of multituberculate mammals before the extinction of dinosaurs. Nature, 483, 457–460.

99. Zhang, X., Shu, D. (2021). Current understanding on the Cambrian Explosion: questions and answers. PalZ, 95, 641–660.

## REFERENCES (for the SI Appendix and Dataset S1

100. Bastide, P., Ané, C., Robin, S., & Mariadassou, M. (2018). Inference of adaptive shifts for multivariate correlated traits. Systematic Biology, 67, 662–680.

101. Bookstein, F. L. (1997). Morphometric tools for landmark data. Cambridge University Press.

102. Clavel, J., Escarguel, G., & Merceron, G. (2015). mvMORPH: an R package for fitting multivariate evolutionary models to morphometric data. Methods in Ecology and Evolution, 6, 1311–1319.

103. Clavel, J., Morlon, H. (2020). Reliable phylogenetic regressions for multivariate comparative data: illustration with the MANOVA and application to the effect of diet on mandible morphology in phyllostomid bats. Systematic biology, 69, 927–943.

104. Cloutier, D., & Thomas, D. W. (1992). Carollia perspicillata. Mammalian Species, 417, 1–9.

105. Cooper, N., Thomas, G. H., Venditti, C., Meade, A., & Freckleton, R. P. (2016). A cautionary note on the use of Ornstein Uhlenbeck models in macroevolutionary studies. Biological Journal of the Linnean Society, 118, 64–77.

106. Fleming, T. H., Hooper, E. T., & Wilson, D. E. (1972). Three Central American bat communities: structure, reproductive cycles, and movement patterns. Ecology, 53, 555–569.

107. Freeman, P. W. (1998). Form, function, and evolution in skulls and teeth of bats; in: Bat Biology and Conservation, T. H. Kunz, Ed. Smithsonian Institution Press, Washington, DC, 140–156.

108. Genoways, H. H., Bickham, J. W., Baker, R. J., & Phillips, C. J. (2005). Bats of Jamaica. Mammalogy Papers: University of Nebraska State Museum, 106.

109. Grossnickle, D. M. (2020). Feeding ecology has a stronger evolutionary influence on functional morphology than on body mass in mammals. Evolution, 74, 610–628.

110. Grossnickle, D. M., Newham, E. (2016). Therian mammals experience an ecomorphological radiation during the Late Cretaceous and selective extinction at the K–Pg boundary. Proceedings of the Royal Society B: Biological Sciences, 283, 20160256.

111. Gual-Suárez, F., & Medellín, R. A. (2021). We eat meat: a review of carnivory in bats. Mammal Review, 51, 540–558.

112. Kushnereit, A. 2004. “Mimon crenulatum” (On-line), Animal Diversity Web. Accessed April 28, 2023 at https://animaldiversity.org/accounts/Mimon_crenulatum/.

113. Kwiecinski, G. G. (2006). Phyllostomus discolor. Mammalian Species, 801, 1–11.

114. López-Aguirre, C., Hand, S. J., Simmons, N. B., Silcox, M. T. (2022). Untangling the ecological signal in the dental morphology in the bat superfamily Noctilionoidea. Journal of Mammalian Evolution, 29, 531– 545.

115. Nowak, R. M. (1999). Walker’s Mammals of the World. John Hopkins University Press. Baltimore, MD.

116. Pellón, J. J., Medina-Espinoza, E. F., Lim, B. K., Cornejo, F., & Medellín, R. A. (2023). Eat what you can, when you can: relatively high arthropod consumption by frugivorous bats in Amazonian Peru. Mammalian Biology, 103, 137–144.

117. Pineda-Munoz, S., & Alroy, J. (2014). Dietary characterization of terrestrial mammals. Proceedings of the Royal Society B: Biological Sciences, 281, 20141173.

118. Quinche, L. L., Santana, S. E., & Rico-Guevara, A. (2022). Morphological specialization to nectarivory in Phyllostomus discolor (Wagner, 1843) (Chiroptera: Phyllostomidae). The Anatomical Record. Early View, doi:10.1002/ar.25147.

119. Reuter, D. M., Hopkins, S. S., & Price, S. A. (2023). What is a mammalian omnivore? Insights into terrestrial mammalian diet diversity, body mass and evolution. Proceedings of the Royal Society B, 290, 20221062.

120. Rojas, D., Ramos Pereira, M. J., Fonseca, C., & Dávalos, L. M. (2018). Eating down the food chain: generalism is not an evolutionary dead end for herbivores. Ecology Letters, 21, 402–410.

121. Rojas, D., Vale, A., Ferrero, V., & Navarro, L. (2011). When did plants become important to leaf-nosed bats? Diversification of feeding habits in the family Phyllostomidae. Molecular Ecology, 20, 22172228.

122. Rojas, D., Vale, A., Ferrero, V., & Navarro, L. (2012). The role of frugivory in the diversification of bats in the Neotropics. Journal of Biogeography, 39, 1948–1960.

123. Selig, K. R., Sargis, E. J., & Silcox, M. T. (2019). Three-dimensional geometric morphometric analysis of treeshrew (Scandentia) lower molars: Insight into dental variation and systematics. The Anatomical Record, 302, 1154–1168.

124. Velazco, P. M. (2005). Morphological phylogeny of the bat genus Platyrrhinus Saussure, 1860 (Chiroptera: Phyllostomidae) with the description of four new species. Fieldiana Zoology, 2005, 1–53.

125. Wetterer, A. L., Rockman, M. V., & Simmons, N. B. (2000). Phylogeny of phyllostomid bats (Mammalia: Chiroptera): data from diverse morphological systems, sex chromosomes, and restriction sites. Bulletin of the American Museum of Natural History, 2000, 1–200.

126. Willig, M. R., Camilo, G. R., & Noble, S. J. (1993). Dietary overlap in frugivorous and insectivorous bats from edaphic cerrado habitats of Brazil. Journal of Mammalogy, 74, 117–128.

127. Wilman, H., Belmaker, J., Simpson, J., de la Rosa, C., Rivadeneira, M. M., & Jetz, W. (2014). EltonTraits 1.0: Species-level foraging attributes of the world’s birds and mammals. Ecology, 95, 2027.

128. Wilson, G. P. (2013). Mammals across the K/Pg boundary in northeastern Montana, USA: dental morphology and body-size patterns reveal extinction selectivity and immigrant-fueled ecospace filling. Paleobiology, 39, 429–469.

129. York, H. A., & Billings, S. A. (2009). Stable-isotope analysis of diets of short-tailed fruit bats (Chiroptera: Phyllostomidae: Carollia). Journal of Mammalogy, 90, 1469–1477.

