## Supporting Information Appendix for "On the cusp of adaptive change: the hierarchical radiation of phyllostomid bats"

### SUPPLEMENTARY RESULTS

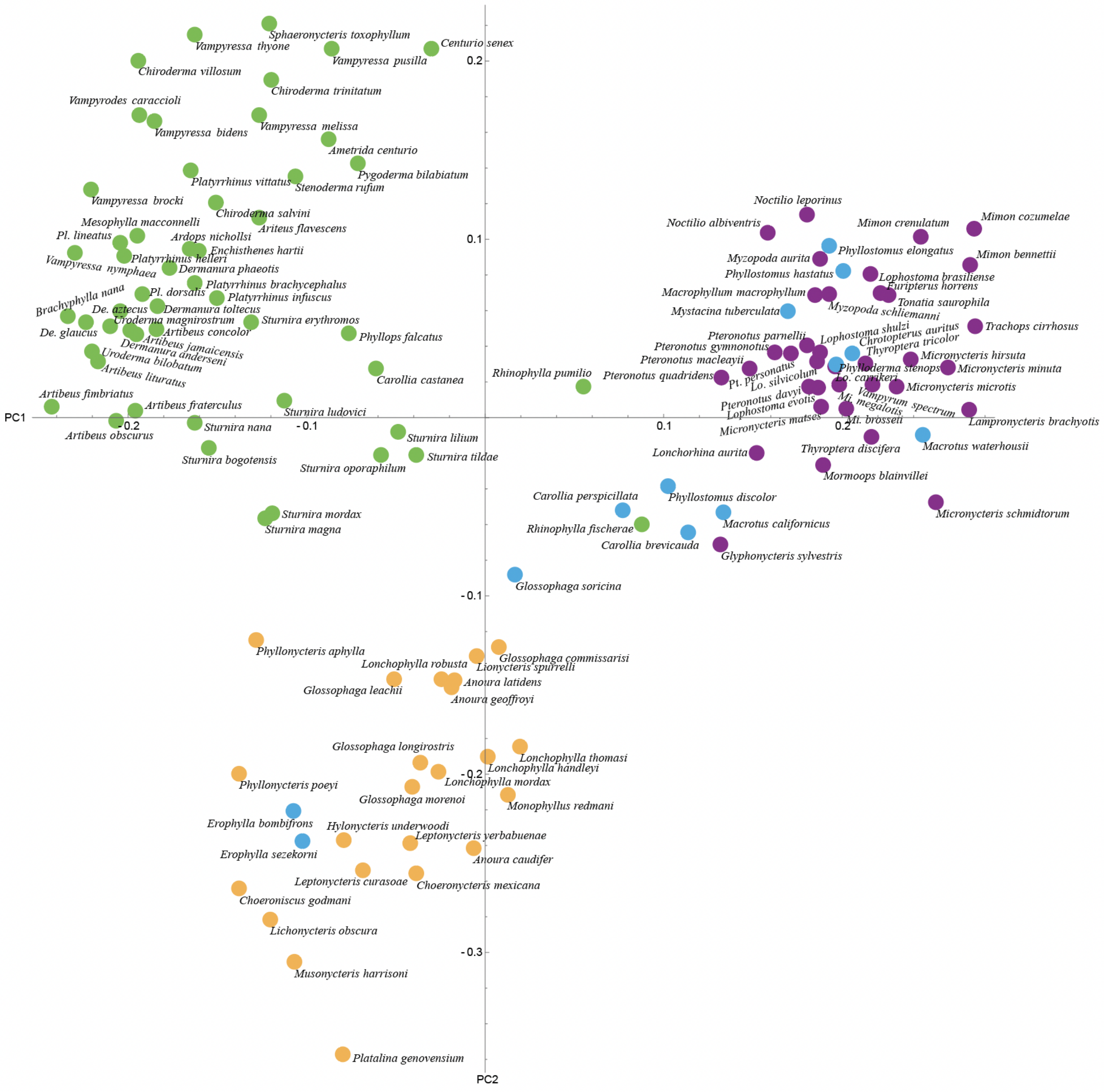


**Fig. S1**. The PC1 and PC2 morphospace plot with species labels. This is the same plot as in Figure 1C, but with labels and without an added phylogeny.

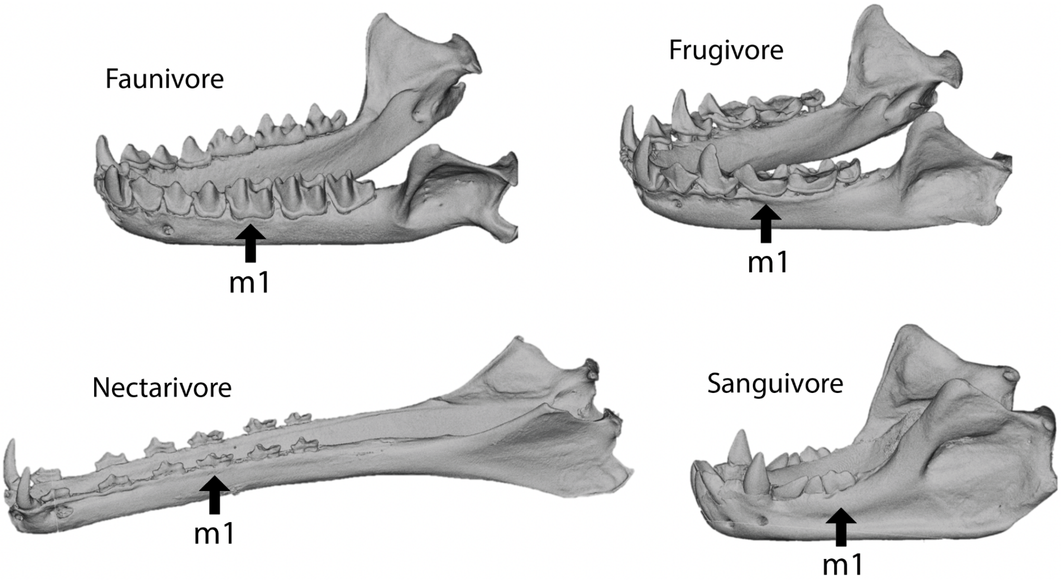


**Fig. S2**. Representative jaws of the major diet groups examined in this study. Sanguivores were excluded due to their derived molars, which are relatively reduced and lack many of the cusps used as landmarks in the geometric morphometrics analyses of molar shape. The jaws are *Lophostoma silvicolum* (faunivore), *Platyrrhinus dorsalis* (frugivore), *Musonycteris harrisoni* (nectarivore), and *Diaemus youngi* (sanguivore).

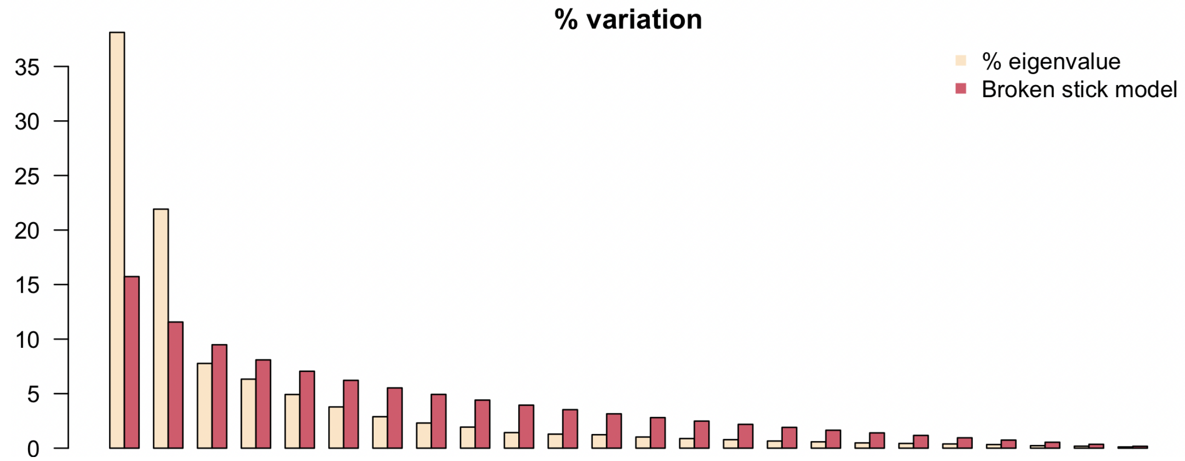


**Fig. S3**. Broken-stick model for testing which principal components (PCs) explain a greater percentage of variance than is expected by chance. Note that the eigenvalues for the first two PCs are considerably greater than the variances of the broken stick model, whereas all other PCs have values that are less than the broken stick model.

**Table S1. Summary statistics for phylogenetic one-way analyses of variance (pANOVAs) and phylogenetic multivariate analyses of variance (pMANOVAs).**

| | | 3 diet categories | | 4 diet categories | |
|---|---|---|---|---|---|
| | | ***F*** | ***p*** | ***F*** | ***p*** |
| pANOVAs | PC1 | 51.9697 | <0.0001 | 37.0092 | <0.0001 |
| | PC2 | 43.4016 | <0.0001 | 21.7008 | <0.0001 |
| | PC3 | 2.1386 | 0.1223 | 1.5101 | 0.2154 |
| | PC4 | 0.9056 | 0.4070 | 0.4351 | 0.7283 |
| | PC5 | 0.2924 | 0.7470 | 1.6613 | 0.1790 |
| | PC6 | 0.4870 | 0.6157 | 1.8252 | 0.1462 |
| | PC7 | 1.2532 | 0.2893 | 1.8573 | 0.1405 |
| | PC8 | 0.2116 | 0.8096 | 0.4833 | 0.6945 |
| | PC9 | 0.1421 | 0.8677 | 0.3734 | 0.7723 |
| | PC10 | 0.2937 | 0.7460 | 0.8164 | 0.4872 |
| | PC11 | 0.1234 | 0.8840 | 0.2079 | 0.8907 |
| | PC12 | 0.3866 | 0.6802 | 0.1143 | 0.9516 |
| | PC13 | 0.9426 | 0.3924 | 1.2664 | 0.2891 |
| | PC14 | 0.6737 | 0.5117 | 0.3300 | 0.8037 |
| | PC15 | 0.0368 | 0.9639 | 0.1809 | 0.9092 |
| | PC16 | 0.9204 | 0.4011 | 1.8870 | 0.1354 |
| | PC17 | 0.0345 | 0.9661 | 0.5551 | 0.6457 |
| | PC18 | 0.4932 | 0.6119 | 0.9826 | 0.4035 |
| | PC19 | 1.2413 | 0.2927 | 0.6851 | 0.5629 |
| | PC20 | 0.0748 | 0.9279 | 0.2389 | 0.8691 |
| | Basicranium width | 1.8987 | 0.1543 | 1.5669 | 0.2011 |
| pMANOVAs | All data, lambda model | 0.0905† | 0.0001 | 0.0729† | 0.0001 |
| | All data, OU model | 0.0326† | 0.0001 | 0.0296† | 0.0001 |
| | PC1–2, lambda model | 0.3143† | <0.0001 | 0.3336† | <0.0001 |
| | PC1–2, EB model | 0.3749† | <0.0001 | 0.3863† | <0.0001 |

See the Methods for descriptions of the three-diet and four-diet categorization schemes. The molar shape principal components analysis (PCA) includes 30 total PCs, but the PCs beyond PC20 often had very small scores that were rounded down to zero by computational programs, and thus we excluded those results here. For pMANOVAs, which were performed using functions in the *mvMORPH R* package (Clavel et al. 2015, Clavel and Morlon 2020), we used the 'lambda' model that transforms phylogenetic branch lengths based on the measured phylogenetic signal. Further, we repeated the pMANOVAs using an EB model for PC1–2 and an OU model for complete molar data (i.e., all Procrustes residuals) because these models were best-fitting to the respective datasets in model-fitting analyses (Fig. 3, Table S3). Note that the inclusion of all Procrustes residuals in pMANOVAs shows weaker molar shape differences among diets compared to the PC1–2-only analysis. *F* statistics are reported for pANOVAs, but for pMANOVAs the test statistics are Wilks lambda, which are marked by the dagger (†).

**Table S2. Results for supplemental measures of disparity (stationary variance) and tests of early burst (EB) patterns.**

| | Stationary variance (from OU1) | | | EB rate decay half-lives (Ma) | | | Node height test (*p* -values) | | |
|---|---|---|---|---|---|---|---|---|---|
| **Sample** | **PC1** | **PC2** | **Body size** | **PC1&2** | **All PCs** | **Body size** | **PC1** | **PC2** | **Body size** |
| Noctilionoidea | 0.152 | 0.064 | 0.056 | 6.098 | inf. | inf. | 0.457 | pos. | pos. |
| Phyllostomidae | 0.191 | 0.097 | 0.078 | 3.690 | inf. | inf. | 0.009 | 0.162 | pos. |
| Phyllostomidae* | 0.166 | 0.109 | 0.037 | 3.382 | 19.524 | inf. | 0.034 | 0.027 | pos. |
| Faunivores | 0.002 | 0.002 | 0.090 | inf. | 2.903*10^8 | inf. | pos. | pos. | pos. |
| Frugivores | 0.016 | 0.008 | 0.024 | 15.961 | 10.827 | inf. | 0.287 | 0.805 | pos. |
| Nectarivores | 0.003 | 0.004 | 0.016 | inf. | 19.452 | inf. | pos. | pos. | pos. |

Stationary variance ($\sigma^2/2\alpha$) is the expected variance when an OU process is stationary, and results are based on the single-regime Ornstein-Uhlenbeck models (OU1) fit to PC1–2. Body size results are based on analyses of basicranium widths, which we used as proxies for body sizes. The Phyllostomidae* clade excludes early-branching phyllostomid faunivores (see red star in Figure 2B). For the node height tests, we only report *p*-values for negative relationships because our goal was to test for an early burst, which is expected to show a negative relationship between node height and morphological contrasts. Abbreviations: EB, early burst; inf., infinity; pos., positive correlation; PC, principal component.

**Table S3. Evolutionary model-fitting results for the 'subgroup analyses.'**

| | | Molar shape | | | | | | Body size | | |
|---|---|---|---|---|---|---|---|---|---|---|
| | | PC1 & 2 (diet correlates) | | | PC1–6 | | | Basicranium width | | |
| **Sample** | **Model** | **ΔAICc** | **weight** | ***r*** | **ΔAICc** | **weight** | ***r*** | **ΔAICc** | **weight** | ***r*** |
| Noctilionoidea | BM1 | 11.603 | 0.003 | – | 59.156 | 0.000 | – | 32.790 | 0.000 | – |
| | OU1 | 26.177 | 0.000 | – | 0 | 1.000 | – | 0 | 1.000 | – |
| | EB | 0.000 | 0.997 | –0.049 | – | – | 0 | – | – | 0 |
| Phyllostomidae | BM1 | 26.499 | 0 | – | 19.109 | 0.000 | – | 0 | 0.788 | – |
| | OU1 | 43.340 | 0 | – | 0 | 1.000 | – | 2.622 | 0.212 | – |
| | EB | 0 | 1.000 | –0.082 | – | – | 0 | – | – | 0 |
| Phyllostomidae* | BM1 | 26.722 | 0 | – | 0.000 | 0.849 | – | 0 | 0.903 | – |
| | OU1 | 45.863 | 0 | – | 4.770 | 0.078 | – | 4.464 | 0.097 | – |
| | EB | 0 | 1.000 | –0.089 | 4.899 | 0.073 | –0.015 | – | – | 0 |
| Faunivores | BM1 | 4.353 | 0.102 | – | 11.432 | 0.003 | – | 30.926 | 0.0000 | – |
| | OU1 | 0 | 0.898 | – | 0 | 0.997 | – | 0 | 1.0000 | – |
| | EB | – | – | 0 | 17.318 | 0.000 | >0.000 | – | – | 0 |
| Frugivores | BM1 | 0 | 0.719 | – | 0 | 0.973 | – | 2.829 | 0.196 | – |
| | OU1 | 12.872 | 0.001 | – | 11.982 | 0.002 | – | 0 | 0.804 | – |
| | EB | 1.885 | 0.280 | –0.019 | 7.359 | 0.025 | –0.028 | – | – | 0 |
| Nectarivores | BM1 | 0 | 0.936 | – | 0 | 0.743 | – | 1.8074 | 0.2883 | – |
| | OU1 | 5.358 | 0.064 | – | 5.358 | 0.051 | – | 0 | 0.7117 | – |
| | EB | – | – | 0 | 2.562 | 0.206 | –0.015 | – | – | 0 |

The Akaike weights (i.e., 'weights') in this table are also plotted in Figure 3. The early burst (EB) model's rate decay parameter (*r*) is used for calculating the rate decay half-lives (Table S2). EB results are omitted in cases where *r* is zero because the model collapses to a Brownian motion (BM) model. See the Methods for more information on the model-fitting analyses and the fitted models.

**Alternative landmarking scheme for frugivores.** The following tables (Tables S4–S6) present results using the alternative landmarking scheme for 20 species of frugivores (see Supplementary Methods). This serves as a sensitivity test for the influence of landmark position uncertainties on our results. The results using the alternative landmarking scheme (Tables S4–S6) are extremely similar to those of the main results (Tables 1, 2, S3), and thus we do not believe that uncertain positions of some landmarks has an influence on our broad conclusions.

**Table S4. Disparity and evolutionary rate analyses using the alternative landmarking scheme for some frugivores.**

| | Disparity (sum of var.) | | MDI | | Rate (from BMM3) | | Stat. variance (OU1) | |
|---|---|---|---|---|---|---|---|---|
| **Sample** | **PC1&2** | **All PCs** | **PC1&2** | **All PCs** | **PC1** | **PC2** | **PC1** | **PC2** |
| Noctilionoidea | 0.043 | 0.070 | –0.155*** | –0.033 | – | – | 0.146 | 0.060 |
| Phyllostomidae | 0.042 | 0.070 | –0.219*** | –0.062** | – | – | 0.196 | 0.083 |
| Phyllostomidae* | 0.039 | 0.068 | –0.230*** | –0.028 | – | – | 0.160 | 0.104 |
| Faunivores | 0.004 | 0.024 | 0.158 | 0.195 | 0.00010 | 0.00010 | 0.002 | 0.002 |
| Frugivores | 0.013 | 0.047 | –0.007 | 0.126 | 0.00035 | 0.00027 | 0.016 | 0.008 |
| Nectarivores | 0.006 | 0.032 | 0.003 | 0.278 | 0.00018 | 0.00029 | 0.003 | 0.004 |

The methodology for these analyses is the same as that for the primary analyses except that alternative paraconid and metaconid landmark positions were used for 20 species of frugivores (see Supplementary Methods). See Methods and Table 1 and S2 captions for more information.

**Table S5. Evolutionary model-fitting results for the 'full sample' analyses using the alternative landmarking scheme.**

| | | PC1&2 (diet correlates) | | PC1–6 | |
|---|---|---|---|---|---|
| | **Model** | **ΔAICc** | **weight** | **ΔAICc** | **weight** |
| **Single-regime** | BM1 | 160.913 | 0.000 | 254.190 | 0.000 |
| | OU1 | 167.274 | 0.000 | 231.657 | 0.000 |
| | EB | 149.755 | 0.000 | 256.417 | 0.000 |
| **Shift** | ***OUBMi*** | 61.805 | 0.000 | 62.234 | 0.000 |
| | OUBM | 68.453 | 0.000 | 71.570 | 0.000 |
| | OUEB | 53.771 | 0.000 | 65.893 | 0.000 |
| | BMEB | 163.017 | 0.000 | 222.392 | 0.000 |
| | BMBM | 108.603 | 0.000 | 111.918 | 0.000 |
| **Multi-regime** | BMM4 | 19.032 | 0.000 | 34.208 | 0.000 |
| | BM1m4 | 55.036 | 0.000 | 153.731 | 0.000 |
| | OUM4 | 58.330 | 0.000 | 97.203 | 0.000 |
| | ***BMM3*** | ***0.000*** | ***1.000*** | ***0.000*** | ***1.000*** |
| | BM1m3 | 52.759 | 0.000 | 147.558 | 0.000 |
| | OUM3 | 45.726 | 0.000 | 108.076 | 0.000 |

The methodology for these analyses is the same as that for the primary analyses except that alternative paraconid and metaconid landmark positions were used for 20 species of frugivores (see Supplementary Methods). See Methods and Table 2 caption for more information.

**Table S6. Evolutionary model-fitting results for the 'subgroup' analyses using the alternative landmarking scheme.**

| | | PC1 & 2 (diet correlates) | | | PC1–6 | | |
|---|---|---|---|---|---|---|---|
| **Sample** | **Model** | **ΔAICc** | **weight** | ***r*** | **ΔAICc** | **weight** | ***r*** |
| Noctilionoidea | BM1 | 11.159 | 0.004 | – | 22.533 | 0.000 | – |
| | OU1 | 25.744 | 0.000 | – | 0 | 1.000 | – |
| | EB | 0 | 0.996 | –0.048 | – | – | 0 |
| Phyllostomidae | BM1 | 25.197 | 0.000 | – | 4.760 | 0.084 | – |
| | OU1 | 42.069 | 0.000 | – | 0 | 0.903 | – |
| | EB | 0 | 1.000 | –0.079 | 8.387 | 0.014 | –0.008 |
| Phyllostomidae* | BM1 | 25.408 | 0.000 | – | 0 | 0.809 | – |
| | OU1 | 44.510 | 0.000 | – | 11.266 | 0.003 | – |
| | EB | 0.000 | 1.000 | –0.087 | 2.912 | 0.189 | –0.030 |
| Faunivores | BM1 | 3.464 | 0.150 | – | 7.517 | 0.023 | – |
| | OU1 | 0 | 0.850 | – | 0 | 0.977 | – |
| | EB | – | – | 0 | – | – | 0 |
| Frugivores | BM1 | 0 | 0.747 | – | 0 | 0.866 | – |
| | OU1 | 12.244 | 0.002 | – | 41.043 | 0.000 | – |
| | EB | 2.180 | 0.251 | –0.008 | 3.737 | 0.134 | –0.015 |
| Nectarivores | BM1 | 0 | 0.974 | – | 0 | 0.995 | – |
| | OU1 | 7.240 | 0.026 | – | 10.726 | 0.005 | – |
| | EB | – | – | 0 | – | – | 0 |

The methodology for these analyses is the same as that for the primary analyses except that alternative paraconid and metaconid landmark positions were used for 20 species of frugivores (see Supplementary Methods). See Methods, Fig. 3 caption, and Table S3 caption for more information.

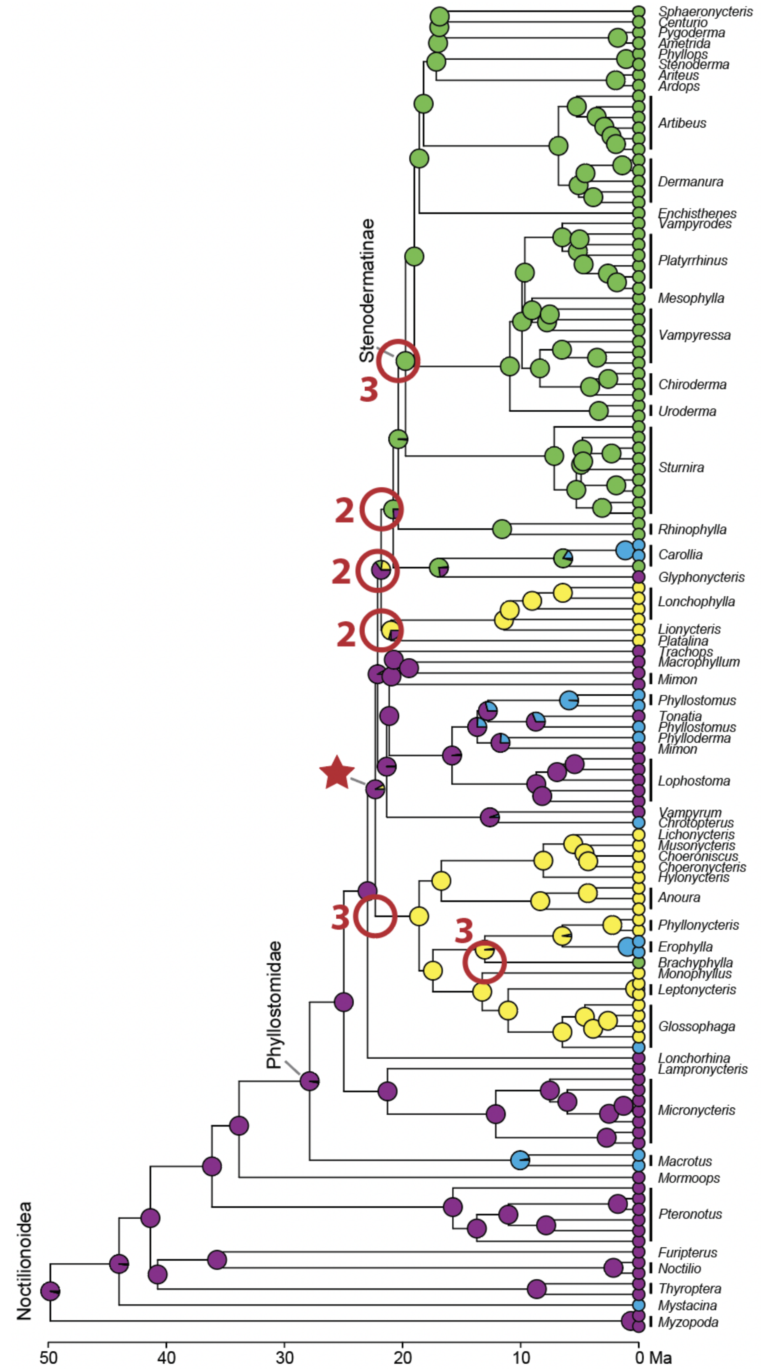


**Fig. S4**. Model-fitting analyses for PC1–2 without *a priori* regime assignments via functions in the *PhylogeneticEM R* package (Bastide et al. 2018). The analysis found evidence of five regime shifts, but there were three equivalent solutions for shift positions, which included six positions in total (marked by red circles). The numbers next to the red circles indicate how many (out of three) equivalent solutions included shifts at those positions. Note that all shifts are within the Phyllostomidae* node and mostly correspond to dietary shifts, consistent with results for the 'full sample' model-fitting analyses with *a priori* regime assignments (Table 2). The *phyloEM* analysis results using PC1–6 indicated 11 regime shifts, but we are not illustrating them here because there were approximately 150 equivalent solutions with various shift locations throughout the tree.

### REFERENCES (for the SI Appendix and Dataset S1)


Bastide, P., Ané, C., Robin, S., & Mariadassou, M. (2018). Inference of adaptive shifts for multivariate correlated traits. *Systematic Biology*, *67*, 662–680.

Bookstein, F. L. (1997). *Morphometric tools for landmark data*. Cambridge University Press.
